# BIO-SPEC: an open-source bench-top parallel bioreactor system

**DOI:** 10.1101/2025.05.21.655256

**Authors:** Laurens Parret, Kenneth Simoens, Jo De Vrieze, Ilse Smets

## Abstract

The BIO-SPEC is an open-source, cost-effective, and modular bench-top bioreactor system designed for batch, sequencing batch, and chemostat cultivation. Featuring thermoelectric condensers to eliminate the need for a chiller, it ensures stable long-term operation. Controlled by a Raspberry Pi, the BIO-SPEC offers flexibility in headplate design, gas supply, and feeding strategies, making it a versatile alternative to high-cost commercial systems. This paper details the design, construction, and validation of the BIO-SPEC system, demonstrating its potential to advance microbiology and bioprocessing research through accessible and reliable hardware at a fraction of the cost.

## 1. Hardware in context

In the field of bioprocessing and biotechnology, laboratory-scale fermentations often rely on flexible and robust bioreactor systems to optimise the process during the research phase [1]. To account for biological variability, replicate reactors are required to run in parallel. By carefully controlling the environmental parameters (dissolved oxygen, pH, temperature, etc.), it is possible to achieve a proxy of a scaled-down fermentation (0.5 - 10 L) that is as realistic as possible. These bench-top systems are constructed from glass or stainless steel, and remain sterilisable by autoclaving, due to their relatively small size. The flexibility in terms of operating conditions (batch/fed-batch/sequencing batch/chemostat) allows for a wide range of applications in microbiology and bio-processing, including for example protein production [2], mammalian cell cultures [3], vaccine development [4], or production of probiotics [5].

As reviewed by Wenzel et al. [6], there is a growing trend in biological research towards the use of do-it-yourself (DIY) lab equipment. This trend is driven by the increasing availability of 3D printing, computer numerical control (CNC) milling, laser cutting, and microcontrollers. In the field of bioprocessing, Wong et al. [7] developed a high-throughput platform (20-40 mL) for microbiological cultivation and growth selection, and Graham et al. [8] as well as Meneses et al. [9] engineered a perfusion flow bioreactor system for cell cultures. Additionally, to reduce the cost of bioreactor operations, Efromson et al. [10] developed a low-cost autosampler, and Jönsson et al. [11] created a 3D-printed peristaltic pump.

Nevertheless, while open-source peripherals are relatively well-developed, the currently available open-source bioreactors are not suitable for axenic, long-term microbial cultivations in the 0.1-1 L scale. For instance, the InnoCell bioreactor [12] is designed for mixed cultures, and therefore not autoclavable. The perfusion systems mentioned earlier [8, 9] are highly relevant, but tailored to their specific application. More versatile initiatives, such as the Biomaker project [13], are dedicated to provide low-level hardware tools which are less relevant in the context considered here. Finally, the Pioreactor project by C. Davidson-Pilon [14] is described in its documentation as a “mostly open-source” bioreactor that provides a flexible platform for microbiological applications. The (commercially available) reactor is essentially a Raspberry-Pi HAT (Hardware Attached on Top) with real-time optical density monitoring, stirring, and temperature control that is designed to fit 20-40 mL (autoclavable) glass vials. For high-throughput screening tests, this is indeed a powerful platform if the proprietary hardware is not an issue. In summary, most open-source bioreactor projects are focused on small-scale, application-specific, or non-sterile use cases using low-voltage electronics.

The high-end standards for commercial bench-top parallel bioreactors of 0.25-10 L are for example the DASGIP^TM^ and DASBOX^TM^ systems of Eppendorf. Also Sartorius offers a competitive alternative with their Biostat^TM^ products. However, while these commercially available systems check all the marks in terms of reliability, ease of use, and technical support, the convenience comes with a high investment cost of 50-150 k EUR depending on the user’s requirements. A more cost-effective DIY alternative that allows long-term cultivation with a similar level of flexibility and longevity of the parts is not yet available. The BIO-SPEC system described in this work aims to bridge that gap by providing a customisable and modular hardware system that is controlled by a Raspberry Pi (RPi). A more elaborate comparison of the BIO-SPEC features with commercial alternatives is provided in supplementary Table S1.

## 2. Hardware description

An overview of the BIO-SPEC system is shown in Figure 1. All of the relevant components are described in more detail in the following sections. The hardware is intended for applications such as:

- microbiological cultures in batch, fed-batch, or continuous (chemostat) operation.
- microbial cultivations where gas supplies and/or feeding strategies need to be varied over time.
- sequencing batch reactors with feast-famine and/or settling regimes in the cycles.
- (bio)chemical reactions in continuously stirred tank reactor (CSTR) operation with (dis-)continuous gas supply.

**Figure 1:**
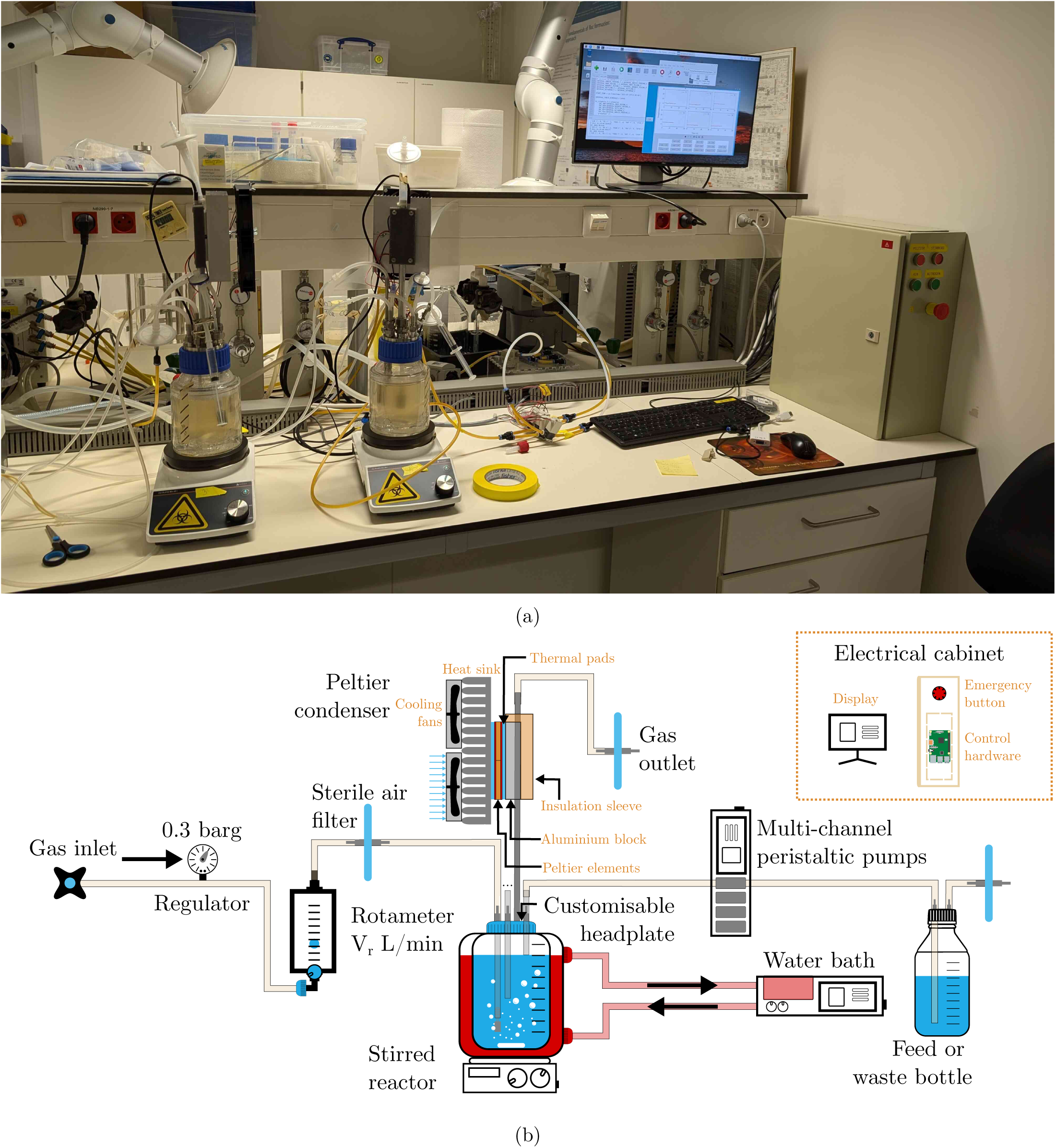
Overview of the BIO-SPEC system. (a) shows a picture of two bioreactors operated in SBR, and (b) indicates a schematic diagram of the relevant connections and components.

### 2.1 Reactor vessel and versatile headplate

The reactor vessel consists of a double-walled (jacketed) glass bottle (250 or 500 mL reactor volume) with a GLS80 screw cap. The broth temperature is then controlled by circulating water through the jacket. If room temperature is sufficient or a climate room is available, however, a wide(r) range of reactor sizes is commercially available at a lower price. A magnetic stirrer bar is added to ensure proper mixing of the culture. If needed, baffles can be inserted to improve mixing, or they can even be permanently integrated by deforming the glass. The latter requires more specialised knowledge, however, and is not necessarily feasible for all users. The content is sealed by a rubber gasket, which is pressed into the glass by the headplate using a screw cap. It is also possible to include a slot in the bottom of the headplate to accommodate an O-ring, which improves handling if the vessel needs to be opened in a laminar flow cabinet. In terms of material requirements, stainless steel (316L) is recommended for its longevity and resistance to aggressive (alkaline) soap, as often used in microbiological research. Aluminium would be a cheaper alternative, but it is less resistant to corrosion and wear.

The headplate is designed to be versatile and customisable. A total of nine ports is suggested for various purposes, as summarised in the list below. This allows for flexibility in the experimental setup and enables researchers to adapt the system to their specific needs. In addition, the headplate can be used with different sizes of reactors, as long as the screw cap is compatible. The proposed headplate design is shown in Figure 2, with the applied ports listed below. The corresponding fittings can either be bought commercially or custom made (see Section 5).

- 2x PG13.5 ports for pH/DO probes
- 3x M12 ports for sampling/withdrawal
- 3x 1/4-28 UNF or M6 ports for feeding/pH control
- 1x M16 port for the off-gas condenser

**Figure 2:**
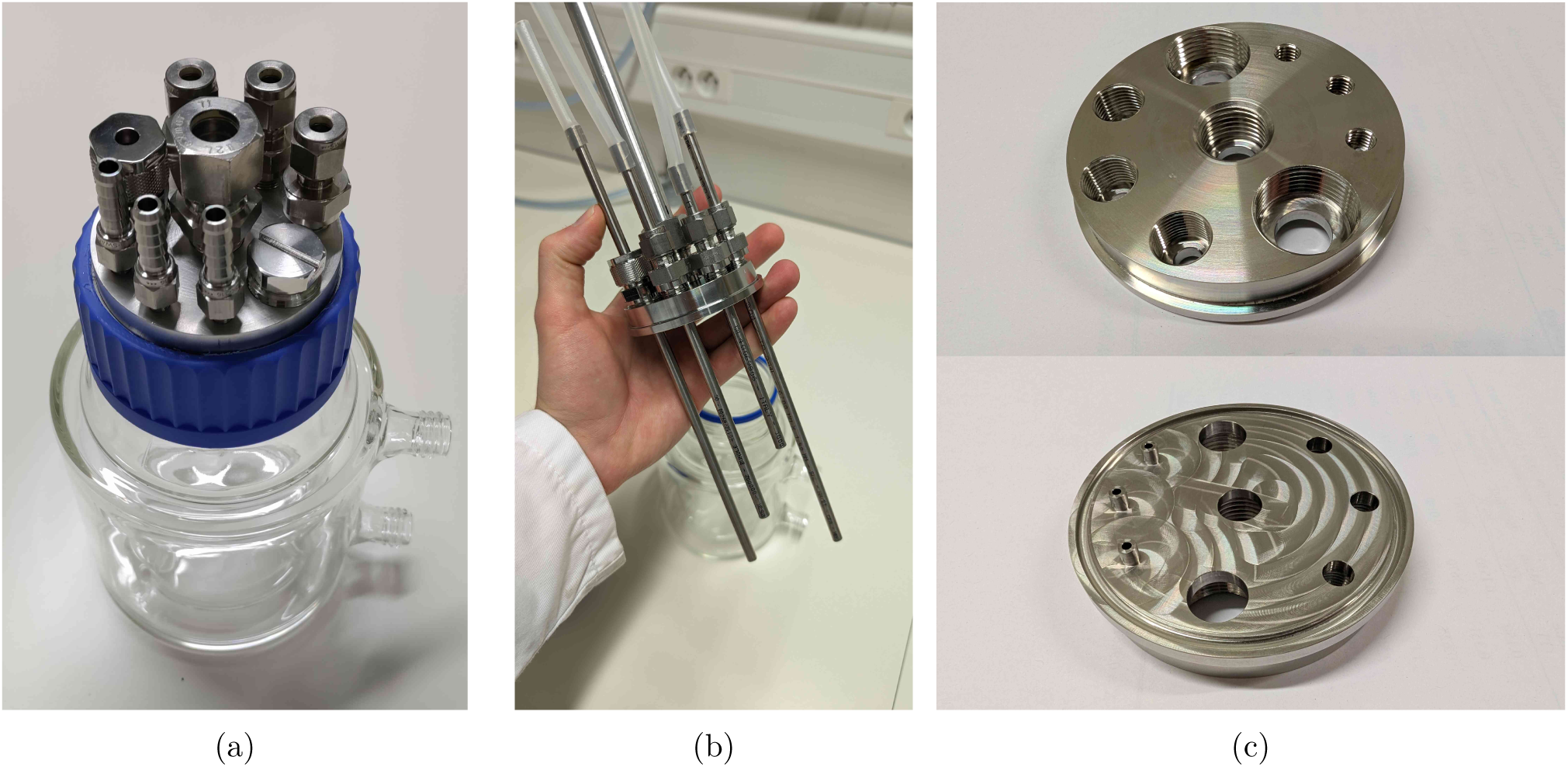
Pictures of the reactor vessel and its headplate design.

Since the assembly is intuitive to customise, the user can choose to add more, fewer, or different types of ports. Evidently, the threads can be adapted to fit other types of (possibly imperial) fittings as well. The only requirement is that the ports are not too close to each other, such that the headplate can easily be assembled. To make the headplate air-tight, the use of O-rings in the blind holes is recommended to reduce wear on the threads due to contact with the reactor content. However, all fittings can also be tightened with PTFE (polytetrafluoroethylene) tape to ensure a proper seal if needed.

Many industry-standard manufacturers offer a wide range of fittings (in terms of threads and sizes) with CAD models available for download, which can be used to verify the compatibility of the custom headplate design. The 1/4-28 UNF thread is especially flexible, since it suits all kinds of fittings, from adapters accommodating PFTE tubing to Luer locks and hose barbs. In many standard labware supplier’s catalogues, such fittings are available in PEEK (polyetheretherketone)/ PP (polypropylene)/stainless steel/…, and allow for straightforward connections to the headplate with minimal space requirements. For example, if clogging and sample tube height are not an issue, no rigid inner tubing is needed and the M12 ports could be replaced by 1/4-28 UNF ports to use PTFE tubing directly for the reactor outlet. The same swap could be interesting if more feeding lines are needed.

As also visible in Figure 2c, the feeding ports have an extra pipe protruding from the headplate in the proposed design. In this way, droplets are formed that directly fall into the reactor liquid. It is possible to leave these out and save on a milling operation, but the feed will start to wet the complete headplate as a result, which may cause corrosion over time. Especially if pH control is desired, the implementation of such pipes is highly recommended. The extra milling step can immediately take care of the slot for an O-ring as well, which is preferred over the gasket. For automated pH control in sterile application, probes with integrated chips can be connected via a COM port on a computer, though to our knowledge, (free) proprietary software is always needed to read them out. If the boundary condition of sterility does not apply, however, cheap pH probes that easily integrate with the RPi are abundantly available.

### 2.2 Thermoelectric condenser

To avoid water loss from the reactor through evaporation, concentrating the broth, and clogging the off-gas filter, a condenser is required. A standard liquid-based condenser requires an external cooling circuit with a chiller that circulates the coolant. This type of solution is not only more expensive, but also not flexible in terms of cooling power and, therefore, easily oversized for its purpose. The BIO-SPEC system uses a Peltier-based condenser shown in Figure 3, which is attached to the off-gas pipe using an aluminium slab and a 3D printed mating block. In this way, the reactor can be autoclaved separately from the condenser unit, and directly mounted on the (aluminium) off-gas pipe when needed. Due to its modular design, cooling is only provided where needed, and the number of reactors can easily be varied between experiments.

**Figure 3:**
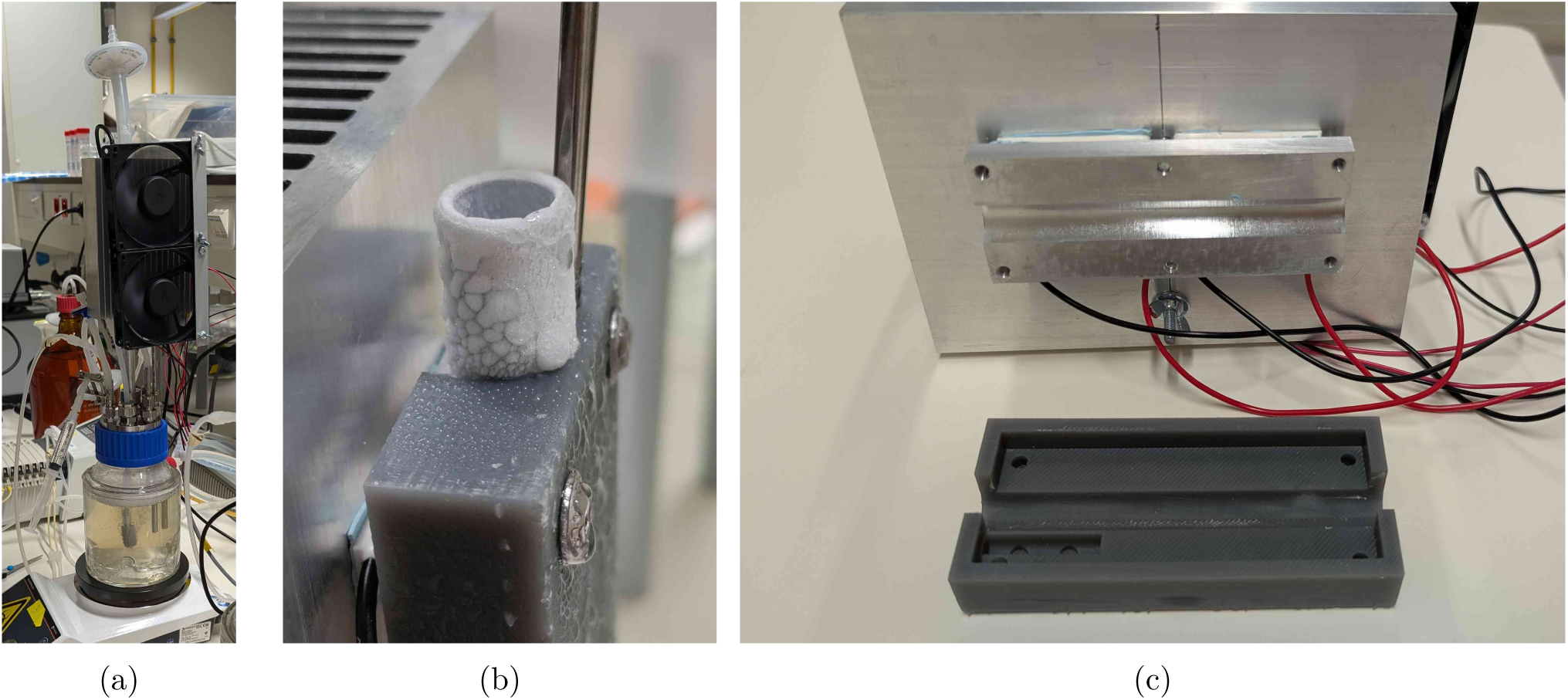
Pictures of the Peltier condenser. (a) shows the condenser in operation, (b) the pipe freezing up without temperature control (red circle), and (c) the slot where the off-gas pipe connects to the condenser components.

The temperature of the aluminium slab is monitored by a DS18B20 temperature sensor connected to the Raspberry Pi (RPi). Each reactor’s thermoelectric coolers and cooling fans are powered by a 12 V power supply, which is controlled by a darlington array. The array turns the power supply on and off based on a controller set to 4 °C. Alternatively, a relay module could be used, though it is larger and more expensive. In any case, without temperature control, the pipe can freeze up (see Figure 3b), potentially causing a pressure build-up in the reactor. Although a PID controller is applied, simple on-off control works well too. The condensed water returns to the reactor vessel, and the off-gas is safely vented through a sterile filter.

As visible in Figure 3, the aluminium heat sink is relatively large, but this is necessary to ensure that the Peltier elements can dissipate the generated heat, even when room temperature rises above *±*25 °C. If room temperature would be controlled, the heat sink can be replaced by a smaller version (see supplementary Figure S1). To facilitate heat transfer, thermal pads are used on both sides of the Peltier elements. In comparison to thermal paste, the pads are easier to apply, and also squeeze slightly when pressure is applied by tightening the nuts during assembly, evening out the heat transfer surface while avoiding fracture of the Peltier elements. The cooling fans are mounted on the heat sink to ensure that the heat is dissipated efficiently, and were selected based on their low noise level and power consumption.

If a chiller (or cooling circuit) that can operate at 4 °C is already available, a coolant-based alternative to the thermoelectric condenser can be built using pipes, tee fittings and bored-through reducers. A schematic diagram using standard Swagelok fittings is suggested in Figure 4. This alternative reduces the complexity of the circuitry and control operations, because no active temperature monitoring and control of the condensers is needed. Since the fittings are to be swaged on the pipes though, the condenser cannot be decoupled from the reactors any more unless plastic ferrules are applied.

**Figure 4:**
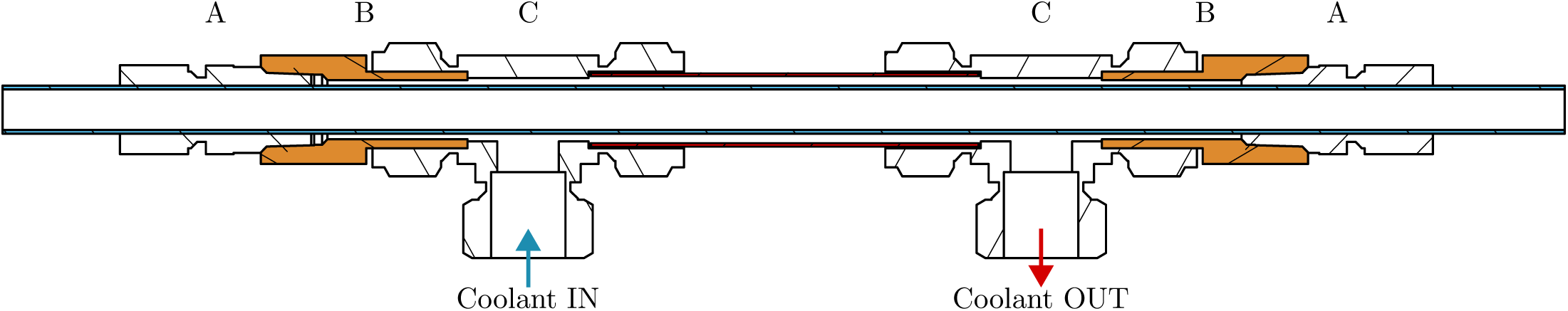
Schematic diagram of a tube-in-tube off-gas condenser based on standard Swagelok parts and their technical datasheets. The required fittings shown here are bored-through reducers (A: 2x SS-12M0-1-8BT), adapters (B: 2x SS-12-TA-7-8), and tees (C: 2x SS-1210-3). The design needs the extra tube adapters (orange, B) to work around the limited bored-through fitting offer for metric pipes (since the off-gas pipe (blue) is 12 mm). If the fittings are bored through by the user, there are not only more size options available, but the adapters can be omitted as well. The outer pipe should have a diameter of 3/4 inch in this example.

In addition to the main BIO-SPEC configuration, a simplified reactor setup was developed for use inside an incubator. This alternative is intended for less demanding applications and eliminates the need for electrical wiring or active temperature control. Instead, it relies on manual operation and a gas humidification system to prevent water loss due to aeration. Evidently, temperature control through double-jacketed vessels is also feasible in that design instead of relying on the incubator temperature. More details on this system and its implementation can be found in Parret et al. [15].

### 2.3 Electrical cabinet & control

All electronics are controlled by a RPi 3 to allow for intuitive programming and more or less plug-and-play GUI integrations in Python. Solenoid valves for gas supply (24 V) and the Peltier condensers (12 V) are controlled using darlington arrays, while stirrers and pumps are controlled by switching the power to DIN-rail sockets (230 V) using a relay board. This choice of components results in a cheap and highly customisable electrical cabinet. Nevertheless, be aware of the fact that 230 V is dangerous, and should be handled with care. Hence, a fuse, an emergency button, proper grounding, and DIN rail terminals for wire connections are mandatory for the high-voltage circuit (see Section 5). The electrical cabinet and its proposed layout is shown in Figure 5. To simplify the high-voltage circuitry, users may opt for 12–24 V peristaltic pumps. In this context, the open-source FAST pump project by Jönsson et al. [11] is a compelling alternative. Although not used in the final experiments here (due to the availability of 230 V peristaltic pumps), the FAST pump was successfully replicated (see Figure 6) and offers a low-cost, RPi-compatible solution for automated feeding, sampling, or waste removal. For more precise control of gas flow, a mass flow controller can be integrated into the aeration system. For example, Bronkhorst mass flow controllers can be directly interfaced with the Raspberry Pi using the open-source ‘Bronkhorst Propar’ Python package, enabling dynamic gas regulation in sensitive applications.

**Figure 5:**
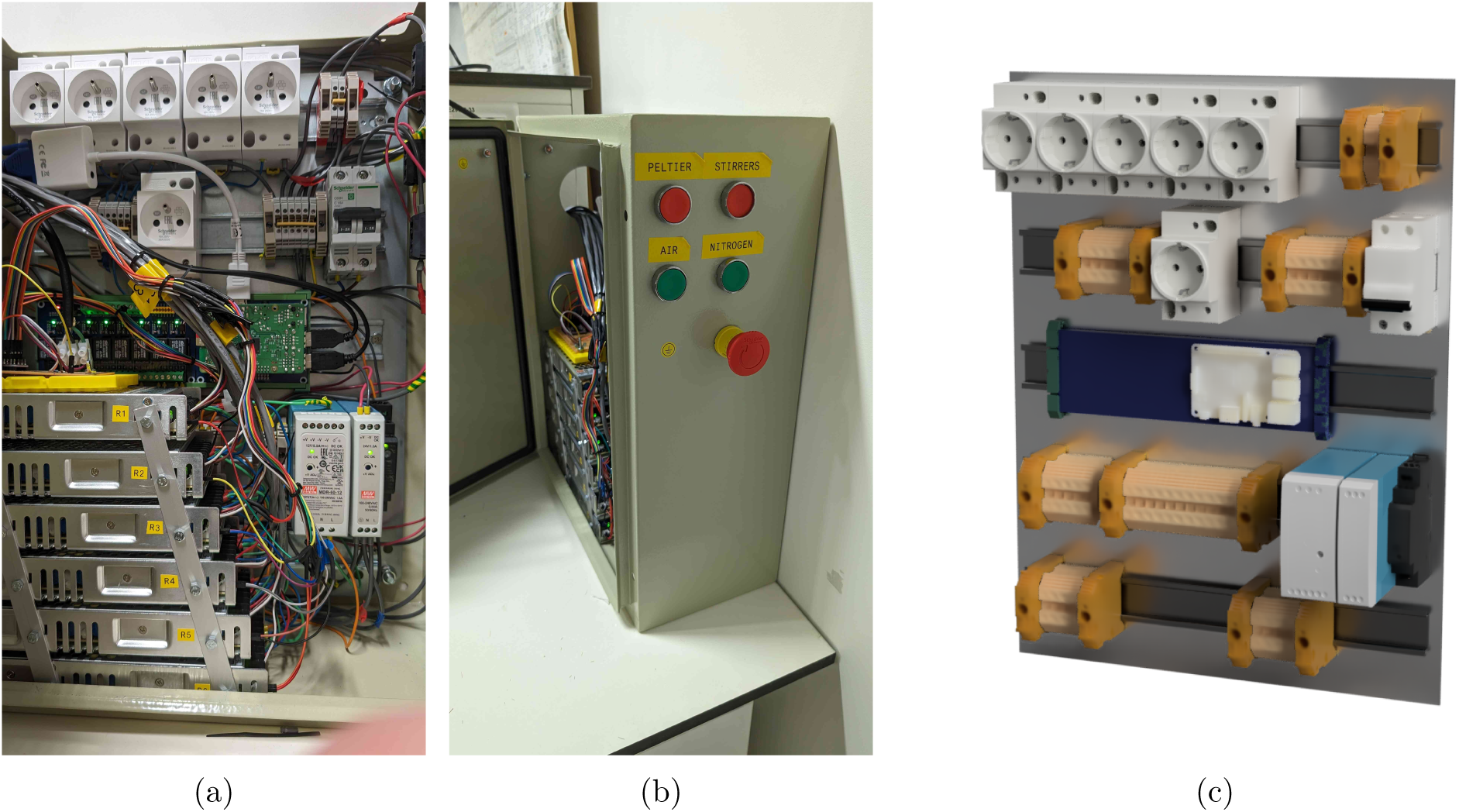
(a,b) pictures of the electrical cabinet, and (c) a render of the component layout. The stacked power supplies shown in (a) power the 12 V thermoelectric condensers. A possible button layout on the side of the cabinet is indicated in (b). The Raspberry Pi can be mounted on the blue DIN rail relay board visible in (c).

**Figure 6:**
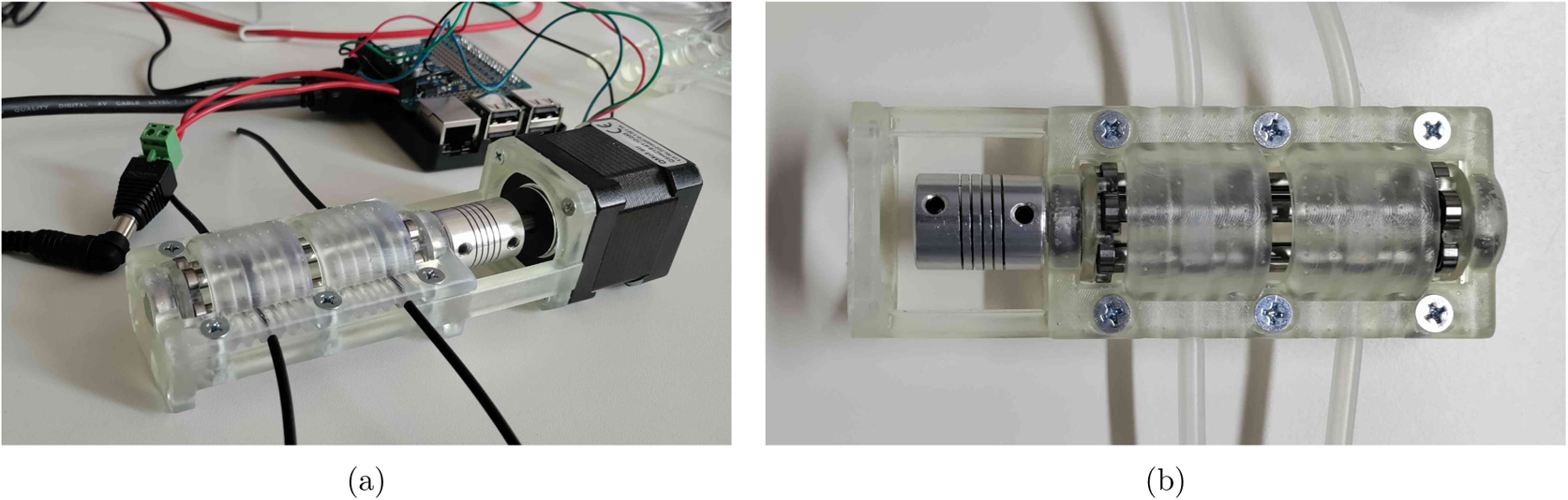
Pictures of the FAST pump as replicated from the design of Jönsson et al. [11].

## 3. Design files summary

All design files associated with the hardware are listed in Table 1. The proposed GLS80 headplate was parametrically modelled in OpenSCAD to allow for intuitive customisation. Threads were included using the ‘threadlib’ library, and can be adapted to other sizes if needed. Do keep in mind that for CNC milling, a .step file is needed, which is why the `Lid_UNF.f3d` (Autodesk Fusion 360, requires the `PG_thread.xml` file to model the PG13.5 threads) and corresponding .step file are included as well. The cap for the GLS80 bottle can be obtained by drilling a 73 mm hole in a commercial one (`GLS80_screw_cap`), or 3D printing the `GLS80_cap.stl` file (possibly in autoclavable resin). The heat transfer plate for the condenser (`aluminium_plate.step`) is designed to be milled from aluminium, with a complementary (3D printed) plastic insulation sleeve `insulation_sleeve.stl`. Electrical wiring schematics are provided in the `schematic_control.pdf` and `schematic_cabinet.pdf` files. A version without the optional buttons is available in the `schematic_cabinet_no_buttons.pdf` file. Two types of custom fittings are provided, one with a Luer lock, and one with a hose barb. To generate the female Luer, the ‘Luer Fitting Generator’ for Autodesk Fusion 360 by Ortus Lab was applied. In case a box to hold the pneumatic components is desired, the `plate*.svg` files can directly be loaded in a lasercutter and assembled. In terms of operating software, the `main_control.py` file is provided, which is written in Python and controls the hardware through the RPi GPIO pins. A second file, `main_control_graph_example.py`, indicates how the code can be extended to the user’s needs by (in this case) plotting the temperature data in real-time. The `requirements.txt` file lists the required Python packages to run the software, while the `tutorial_main_control.html` file pro-vides an elaborate explanation of the code, the implementation decisions, and how the script can be adapted to the user requirements.

**Table 1:**
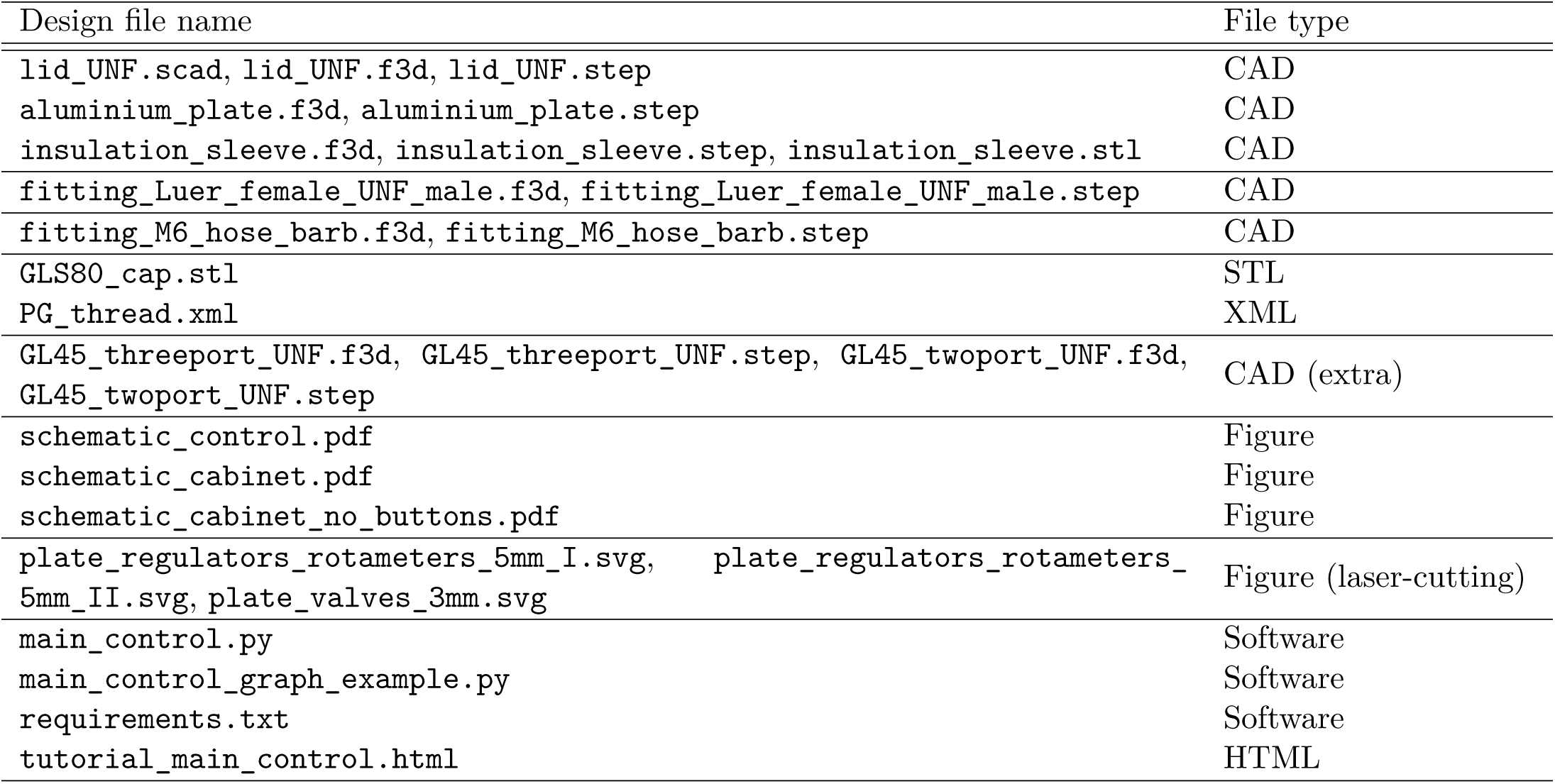
Summary of design files for the BIO-SPEC system. All design files are available at https://doi.org/10.17605/OSF.IO/39WSB, with the open-source license GNU GPL v3.

| Design file name | File type |
| --- | --- |
| `lid_UNF.scad`, `lid_UNF.f3d`, `lid_UNF.step` | CAD |
| `aluminium_plate.f3d`, `aluminium_plate.step` | CAD |
| `insulation_sleeve.f3d`, `insulation_sleeve.step`, `insulation_sleeve.stl` | CAD |
| `fitting_Luer_female_UNF_male.f3d`, `fitting_Luer_female_UNF_male.step` | CAD |
| `fitting_M6_hose_barb.f3d`, `fitting_M6_hose_barb.step` | CAD |
| `GLS80_cap.stl` | STL |
| `PG_thread.xml` | XML |
| `GL45_threeport_UNF.f3d`, `GL45_threeport_UNF.step`, `GL45_twoport_UNF.f3d`, `GL45_twoport_UNF.step` | CAD (extra) |
| `schematic_control.pdf` | Figure |
| `schematic_cabinet.pdf` | Figure |
| `schematic_cabinet_no_buttons.pdf` | Figure |
| `plate_regulators_rotameters_5mm_I.svg`, `plate_regulators_rotameters_5mm_II.svg`, `plate_valves_3mm.svg` | Figure (laser-cutting) |
| `main_control.py` | Software |
| `main_control_graph_example.py` | Software |
| `requirements.txt` | Software |
| `tutorial_main_control.html` | HTML |

## 4. Bill of materials summary

The complete bill of materials can be found in the `bill_of_materials.xlsx` file (https://doi.org/10.17605/ OSF.IO/39WSB), alongside more information on where to source the components. Both a suggested vendor as well as the corresponding part number are provided for every component `’designator’` that is mentioned in this text. A summary of the cost breakdown for the bill of materials is provided in Table 2. Overall, the largest fixed cost is associated with the electrical cabinet, while the fittings and reactor vessel are the main contributors to the variable cost. The total fixed and variable costs are approximately 1610 EUR and 1110 EUR, respectively. This means that the system can be scaled up to multiple reactors at a relatively low cost per additional reactor.

**Table 2:** Overview of the estimated costs for the BIO-SPEC system components as required for a single controlled reactor vessel. The costs are categorised by the function of the components, with approximate values provided in euros. Fixed costs cover the entire system, while variable costs are required for every reactor. The values shown here are indicative and may vary based on supplier and order quantity.

| Label | Total cost | Fixed cost | Variable cost |
| --- | --- | --- | --- |
| Condenser | 220 | 130 | 90 |
| Electrical cabinet | 890 | 880 | 10 |
| Fittings | 330 | 90 | 240 |
| Gas preparation | 310 | 140 | 160 |
| General | 100 | 100 |  |
| Reactor vessel | 680 | 200 | 470 |
| Tools | 80 | 80 |  |
| Custom | 120 |  | 120 |
| Grand total | 2720 | 1610 | 1110 |

## 5. Build instructions

### 5.1 Reactor headplate and vessel

As already mentioned in Section 2, the headplate (`lid_UNF.step`) needs to be (CNC) manufactured out of (preferably) stainless steel 316L. It is also possible to 3D print the part, though for precision and autoclavability, it is necessary to use a high-quality resin and printer to ensure that the threads come out nicely. As a proof-of-concept, the headplate was successfully printed in Formlabs BioMed Amber Resin^TM^, as shown in Figure 7c. In this case, it is recommended to implement a larger tolerance for the threads to loosen the fit and limit wear on the plastic. Regardless of the material, the threads can be improved afterwards using the respective taps (e.g. `PG13.5_tap`) if needed. Figures 7a and 7b show the stainless steel version with appropriate O rings (`O_ring_UNF/M12/M16/Lid`) in the blind holes and bottom groove. The milling toolpath for the bottom of the headplate may remain visible if extra polishing steps are left out, as seen in Figure 7b.

**Figure 7:**
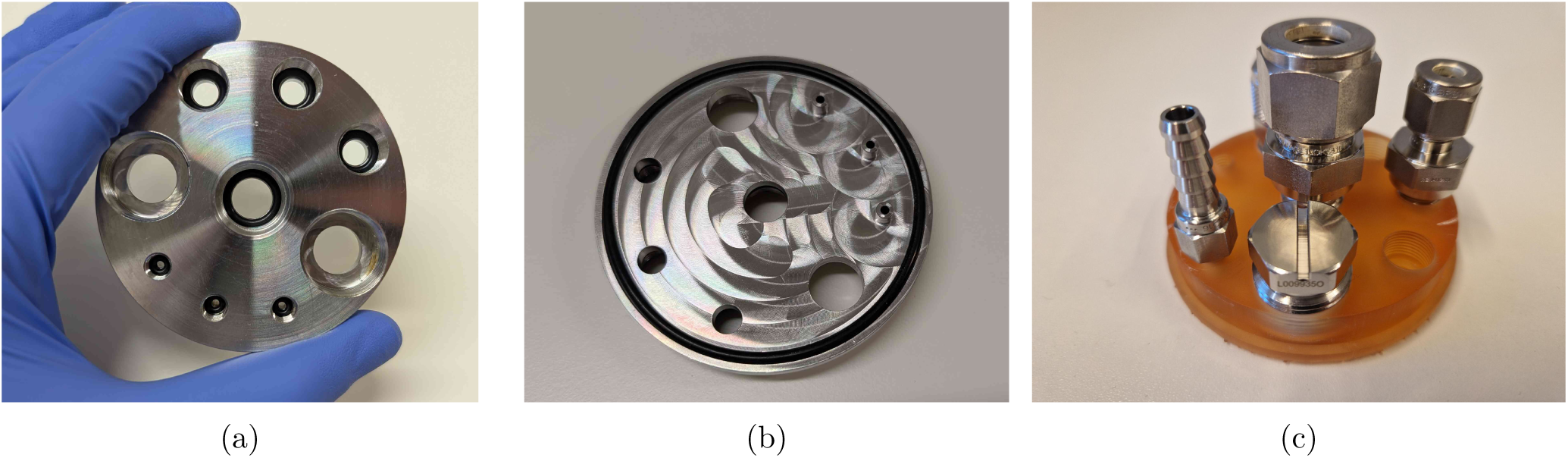
Pictures of the manufactured headplates. (a) and (b) show the stainless steel version with appropriate O rings in the blind holes, while (c) shows the 3D printed version in Formlabs BioMed Amber Resin^TM^.

The complementary fittings need to be manufactured out of stainless steel as well (e.g. `fitting_Luer_ female_UNF_male.step`), or purchased in stainless steel or polypropylene (1x `PG13.5_6_mm_fitting_Eppendorf`, 1x `blind_plug_PG13.5_Eppendorf`, 3x `M12_6_mm_fitting_Swagelok`, 1x `M16_12_mm_fitting_gas_outlet`, 2x `UNF_barb_fitting_Masterflex` for each reactor). If you would opt to swap out the UNF threads with M6, a hose-barb alternative (`fitting_M6_hose_barb.step`) can be manufactured instead (also shown in Figure 2a).

1. Cut the metal tubes (`6_mm_SS_tube`) to the desired length. A tube cutter can in principle be used, but this slightly deforms the tube at the end, making the use of metal ferrules impossible. Just cutting it with a saw and filing the ends down is therefore recommended if the tubes are to be (permanently) swaged at a specific height. The suggested tube has an inner diameter of 3 mm to limit the dead volume in the tubing, while still avoiding clogging due to biomass.
2. Drill a 6.5 mm hole (using `drill_bit_for_6_mm_Swagelok_fittings`) in the `M12_6_mm_fitting_Swagelok` tubes to ensure the pipes fit through. Make sure to use something similar to the `cutting_oil` compound when drilling through stainless steel. Alternatively, bored-through fittings can be bought directly from Swagelok as well and require no drilling operations.
3. Cut the aluminium off-gas tube (`off_gas_pipe_aluminium`) to length (at least 250 mm), insert it into the `M16_12_mm_fitting_gas_outlet` compression fitting and permanently fix it in place by tightening the nut with a wrench according to the manufacturer’s instructions. It is possible to use a different size for the off-gas tube, but keep in mind that the inner diameter needs to be large enough for the condensed droplets to easily drip back into the reactor. If the tube is too narrow, water will be entrained in the gas flow, and the condenser will not work as intended. In addition, be aware that a lot of cheap aluminium pipes are sold at general DIY stores with wide tolerances in the diameter. These pipes do not comply with the specifications of the compression fitting, and would need to be turned to size.
4. Insert all fittings into the headplate containing the O-rings, and tighten by hand (Figure 8a). Probes with PG13.5 thread can be inserted as well, possibly after prior calibration.

a. Insert the metal tubes or PTFE (for aeration, `PTFE_tubing_gases`) into the fittings, and slip over the ferrules, backferrules, and O-rings, as indicated in Figure 8b for the aeration fitting. Adding the aeration stone (`aeration_stone`) and PTFE aeration filter (`sterile_filter_gas`) is done by slipping silicone tubing over the PTFE tube, and securing the connection with autoclavable cable ties (`zip_tie_polypropylene`) as indicated in Figure 8c.
b. Adjust the height of the tubes by pouring the desired amount of water in the vessel (`GLS80_vessel`). In the example shown in Figure 9, one tube is applied for withdrawal at the 350 mL mark, while the other is only there for safety reasons, slightly above the maximum feed level of 400 mL. The outlet of the safety tube is then connected in parallel and counter-current with the feeding line on the same pump, to avoid overfilling the reactor if withdrawal was not completed due to a pump malfunction. In any case, it is advised to keep at least 20% headspace inside the vessel. The height of the tubes can be adjusted by loosening the fitting, and sliding the tube up or down. At the desired height, the fitting can either be permanently swaged, or just tightened when using plastic ferrules. The polypropylene Eppendorf ferrules work equally well on the 6 mm metal tubes.
c. Add silicone tubing to all outer connections, insert female Luer lock fittings `female_Luer_lock_ fitting_1/4` and cover in aluminium foil for autoclaving. The off-gas PTFE filter is also attached using silicone tubing (`tubing_gas_outlet`) and PP zip ties. All corresponding tubing are autoclaved separately with male Luer fittings (`male_Luer_lock_fitting_1/4`).
d. A sampling valve (`sampling_valve_Luer`) can be added to both the sampling pipe and the custom `fitting_Luer_female_UNF_male` fitting. Combining the latter with such a sampling valve is useful to inoculate the vessel. By using a blunt needle and a syringe, the valve acts as a kind of septum, allowing the aseptic transfer of the inoculum (see Section 6.2). The final reactor headplate assembly is shown in Figure 9c. Lastly, medium can be added through the sampling valve after autoclaving (possibly through a filter if needed). Alternatively, the vessel can be opened in a biosafety cabinet to aseptically transfer medium to the reactors.

**Figure 8:**
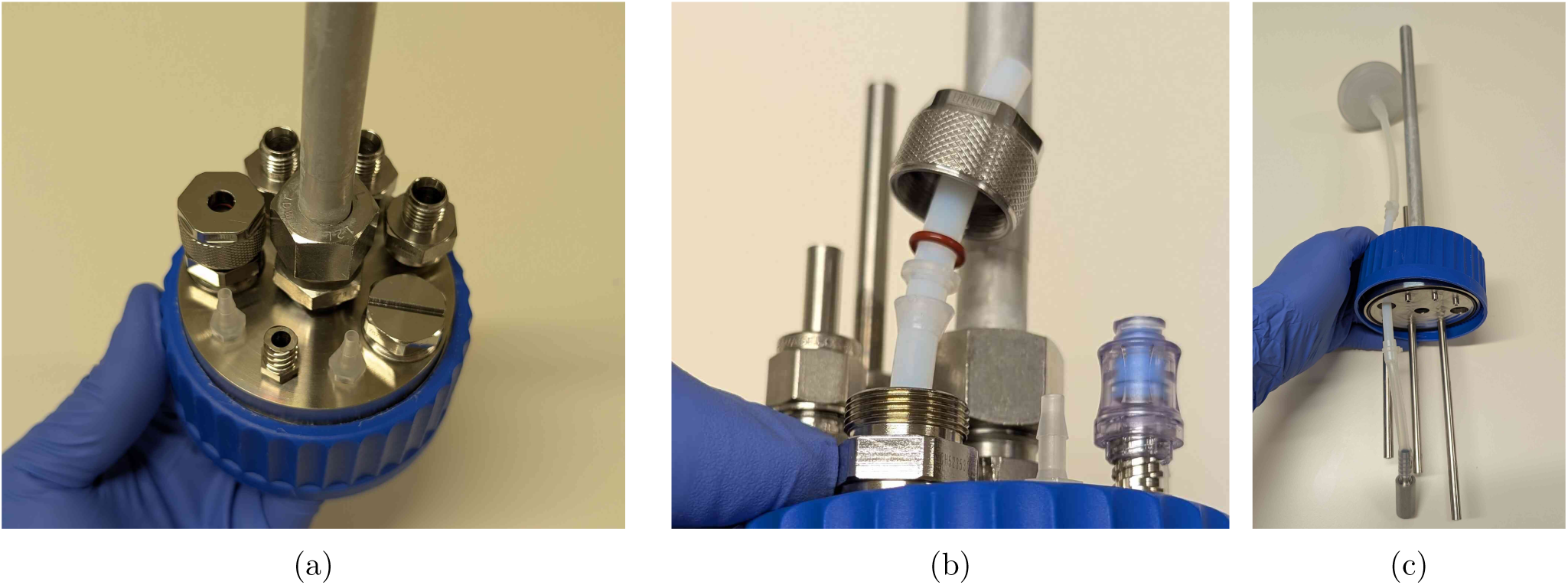
Assembly of the reactor lid, fittings, and aeration.

**Figure 9:**
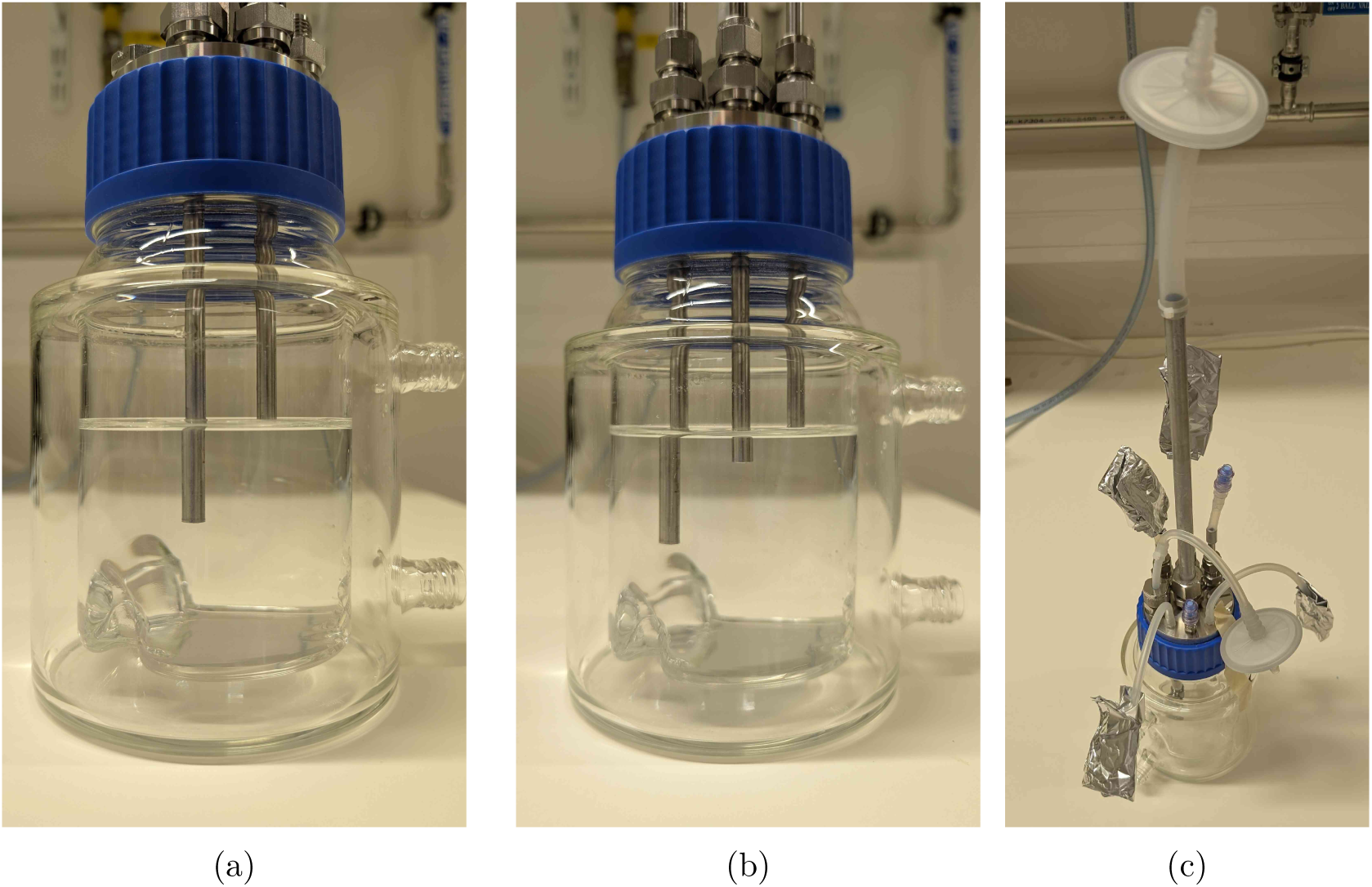
Reactor level calibration using tube heights (a-b) and the final reactor headplate assembly (c).

### 5.2 Thermoelectric condenser for each reactor

1. Mill the aluminium plate `aluminium_plate.step` and make sure the holes are tapped to fit M4 screws.
2. Print the insulation sleeve (`insulation_sleeve.stl`). Here, Formlabs Draft Resin^TM^ was used (Figure 11a), but any other printer or plastic should be suitable for this part as well. The sleeve is designed to fit the 12 mm off-gas tube and the temperature probe. Both are then tightly pressed against the aluminium plate using M4 screws (`M4_screw_16mm`). This does not only ensure good heat transfer, but also keeps the condenser in place.
3. Drill three 4.5 mm holes in the middle of the heat sink (using `drill_bit_for_condenser`) according to the drawing in Figure 10a. The holes to accommodate the aluminium plate are 32 mm apart. Try to drill in between the fins, at high rpm and with minimal downward pressure to avoid damaging the drill bit (and the heat sink). The location of the third hole to hold the fans is not that critical.
4. Assemble the thermoelectric coolers (2x `peltier_element`), aluminium plate, and heat sink (`aluminium_ heat_sink`) as shown in Figures 10b and 10c. Ensure the cold side of the thermoelectric cooler is facing the aluminium plate (this can be double-checked easily by connecting it to a battery). Place a thermal pad (`thermal_pad`) between the Peltier elements and the heat sink, as well as between the Peltier elements and the aluminium plate (Figure 10b). Two long M4 bolts (`M4_bolt_50mm`) press the assembly together, but should not protrude out of the aluminium plate. If needed, extra washers (`M4_washer`) can be added to ensure the screws do not touch the insulation sleeve (as also visible in Figure 10c). Tighten the bolts alternately until the thermal pads are compressed, but do not overtighten to avoid shattering the Peltier elements.
5. The fans (2x `fan`) are mounted on the hot side of the Peltier elements to improve heat dissipation. Align a piece of aluminium (`aluminium_bar`) with the mounting holes of the fans, and drill accordingly. The fans are then attached to the rail using M4 bolts and wing nuts (`M4_bolt_30mm`, `M4_hex_nut`) as shown in Figure 10c. The rail is fixed to the heat sink using a piece of M4 threaded rod (`M4_rod`), hex nuts (`M4_hex_nut`) and washers (`M4_washer`) (Figure 11b and 11c), adjusting the height to keep the fans a maximum of 1 cm above the heat sink. Tighten the wing nuts on the rod to ensure the fans are securely attached to the heat sink.
6. Every complete condenser can be attached to the off-gas pipe of the reactor, as shown in Figure 11. The insulation sleeve should be placed around the off-gas pipe, and the condenser assembly is then fixed in place using the hex screws (4x `M4_screw_16mm`). The temperature probe (`temperature_sensor`) is inserted in the insulation sleeve before fully tightening the assembly.

### 5.3 Electrical cabinet (high-voltage)

The suggested electrical cabinet (`cabinet`) tightly fits all of the required components. If you wish to extend or modify the setup later on, it is recommended to buy a larger version. As already stressed earlier, in the steps provided below, you will need to make connections for a 230 V circuit to control magnetic stirrers and 230 V pumps. Always be mindful of proper grounding and insulation to avoid electrical hazards. For the high-voltage circuit, ensure the wiring is appropriately sized (14-12 AWG, e.g., `wire_14_AWG`) as well.

1. Drill holes in the electrical cabinet to accommodate the desired button layout (`hole_saw_22mm`). If the cabinet needs to remain IP66, all cables should be fed through the bottom of the cabinet, and sealed with a cable gland. Here, instead, a large hole was cut in the side of the cabinet (`hole_saw_73mm`) to allow for easy access.

- For the sake of heat transfer, it is also recommended to cut out space for extra cooling fans in the side. In the current design, only two fans are added at the top, leading to potential overheating of the Peltier power supplies at high ambient temperatures.
- After cutting through the steel cabinet, make sure to sand down the edges (e.g. using sandpaper or a Dremel^TM^) to avoid cutting hazards on the burrs. Be aware that a lot of sparks will form when cutting or deburring this material, so take appropriate safety measures and wear personal protective equipment (PPE).
- Insert the buttons (`emergency_button` and for example `latching_push_button_red`) according to the manufacturer’s instructions.
2. Attach five DIN rails (`DIN_rail`) to the metal sheet of the cabinet using M4 screws (`M4_bolt_10mm`).
3. Stack the power supplies for the Peltier elements (`power_supply_peltier`) using short M4 screws (`M4_ bolt_10mm`) and a piece of aluminium rail. Ensure there is enough space between the power supplies to allow for proper cooling. The rack should then fit inside the cabinet as shown in Figure 12a.
  - Connect all power supplies of the rack in parallel on their GND, live and neutral lines, and also add three extra wires to a terminal strip (`terminal_strip_3_way`). The latter can be then be connected to the main power line later on. In this way, you can easily detach the rack from the cabinet without having to disconnect every individual power supply.
4. Mount the following components to the DIN rails as indicated in the proposed layout of Figures 12 and 14: 1x `relay_module`, 1x `power_supply_24V`, 1x `power_supply_12V`, 1x `power_supply_5V`, 1x `circuit_ breaker`, and 6x `DIN_socket`.
5. Connect the live and neutral wires of a plug to the circuit breaker, and the ground wire to a grounding terminal block. Pass the live wire from the breaker through the emergency button before connecting it to the rest of the circuit.
6. Ground the cabinet and its door by connecting it to the earth wire of the power line as detailed in the manufacturer’s instructions.
7. Connect all of the components according to the design file `schematic_cabinet.pdf` using terminal blocks (`DIN_rail_terminal_block`) and (preferably) bootlace ferrules (e.g. `wire_management_ferrule`) at the wire ends. The latter avoids stray copper wires and an easier/cleaner connection overall. Allow appropriate spacing between arrays of live, GND and neutral blocks, and keep them tidy using end brackets (`DIN_rail_ end_bracket`). The terminal blocks can be interconnected at the front using a jumper bar (`jumper_bar`, cut to size). To avoid wire clutter, also consider using zip ties and mounts (`zip_tie,cable_tie_mount`). Examples of the required connections are shown in Figure 13. If you want to include optional manual control buttons, a suggested layout is provided in the design file `schematic_cabinet_with_buttons.pdf`. The layout can be adapted to your needs.
8. Mount the RPi to the relay board. Consult the manufacturer’s instructions for the correct GPIO pin layout to control the relays to test the setup in its current form. Double-check all connections before plugging in the main power line and switching on the circuit breaker. Note that the RPi is directly powered by the 5 V connection to the relay board.
9. Attach two fans to the top of the cabinet (using zip ties) to improve cooling of the power supplies. They can be permanently connected to the 12 V line.

**Figure 10:**
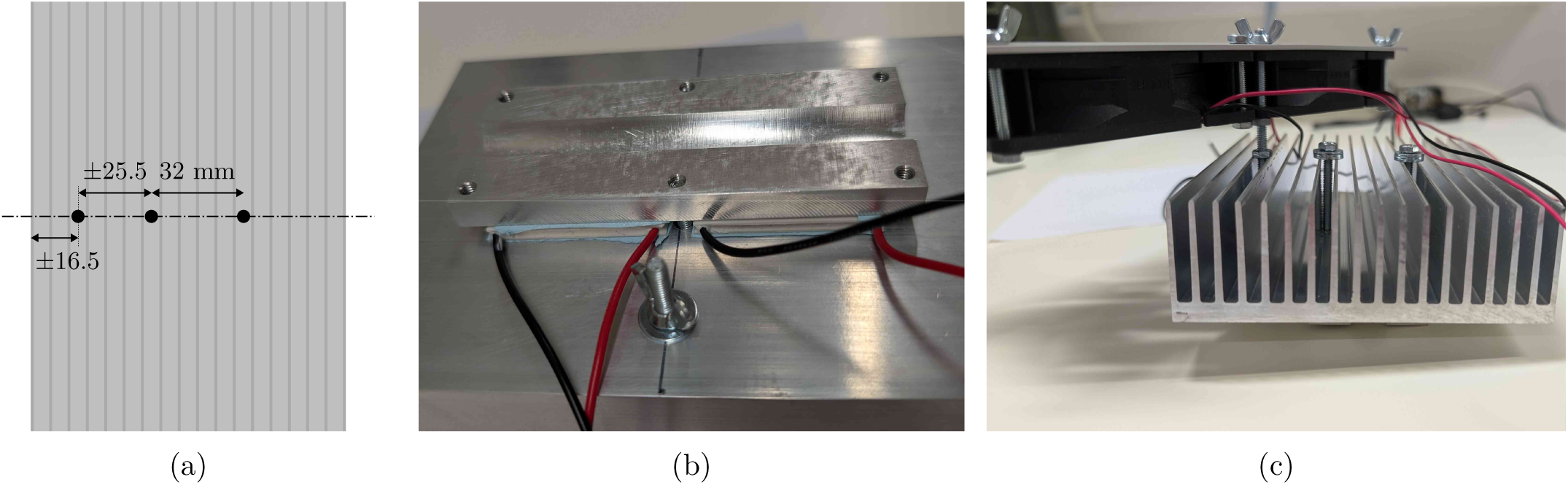
Heat sink assembly. (a) indicates where to drill the holes in the heat sink, (b) the assembly at the hot side of the Peltier elements, and (c) the assembly at the cold side.

**Figure 11:**
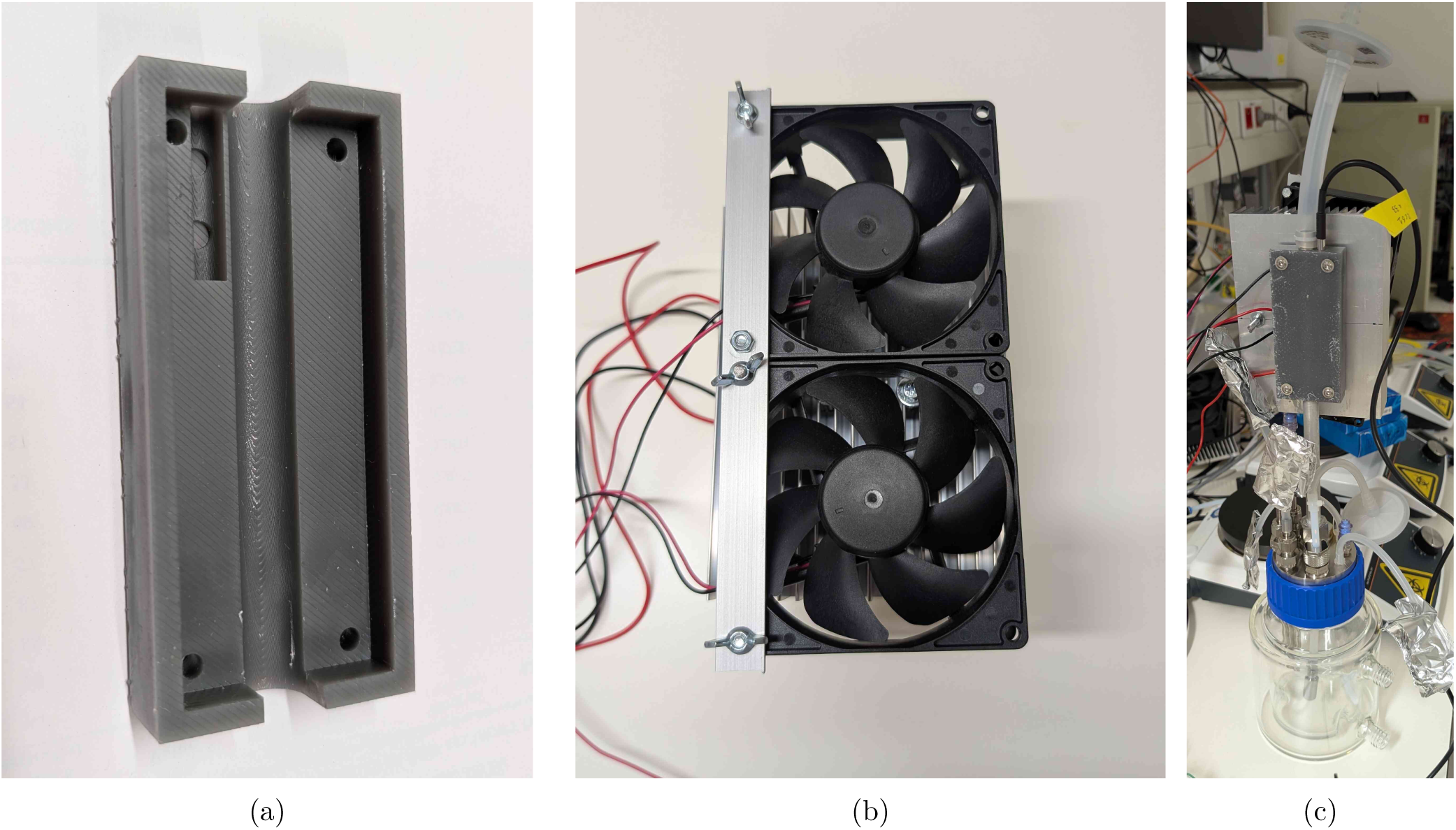
Thermoelectric condenser assembly. (a) shows the insulation sleeve, (b) the assembly of the Peltier elements, aluminium plate, and heat sink, and (c) the final assembly on the off-gas pipe.

**Figure 12:**
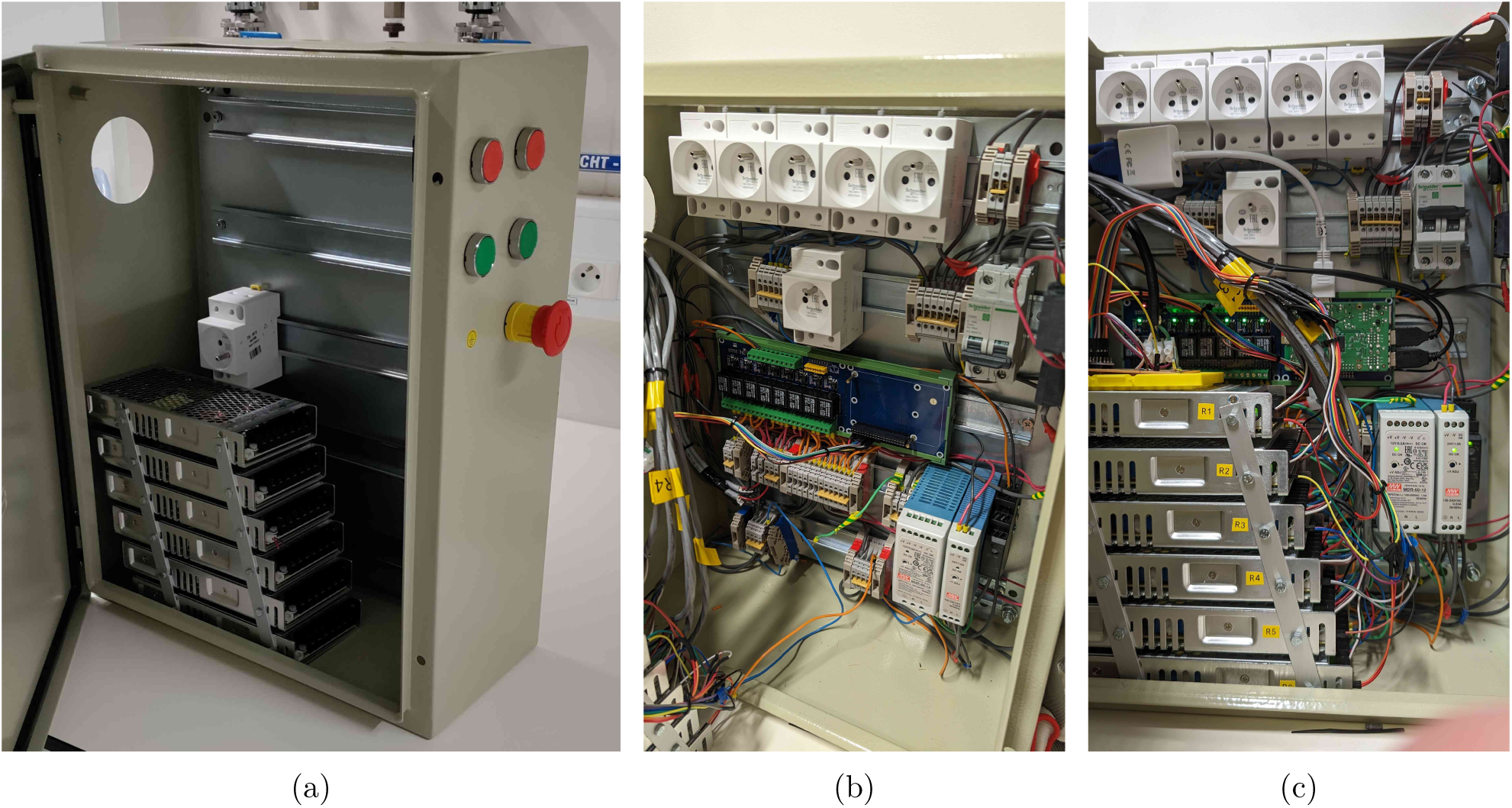
Electrical cabinet assembly. (a) shows the bare cabinet with the power supply rack, while (b) and (c) show the cabinet with all components mounted. A 3D render of the cabinet layout is provided in Figure 14.

**Figure 13:**
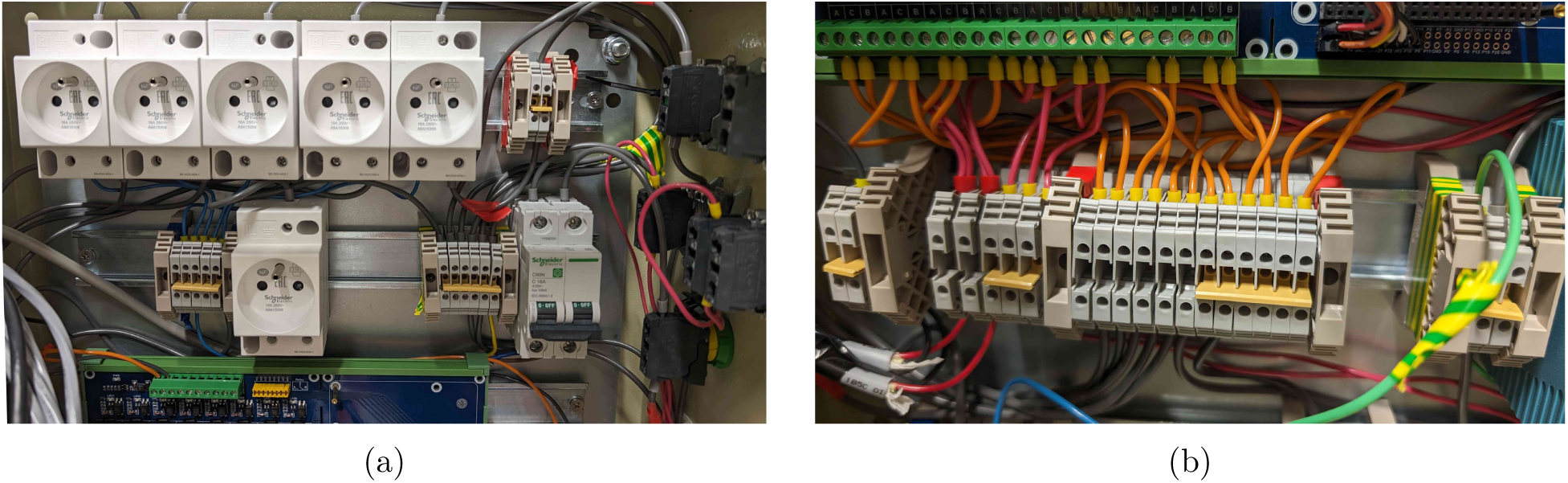
Example close-up pictures of electrical connections using terminal blocks. Arrays of live, neutral and ground terminal blocks are labelled with red, blue and green-yellow electrical tape at the end brackets.

**Figure 14:**
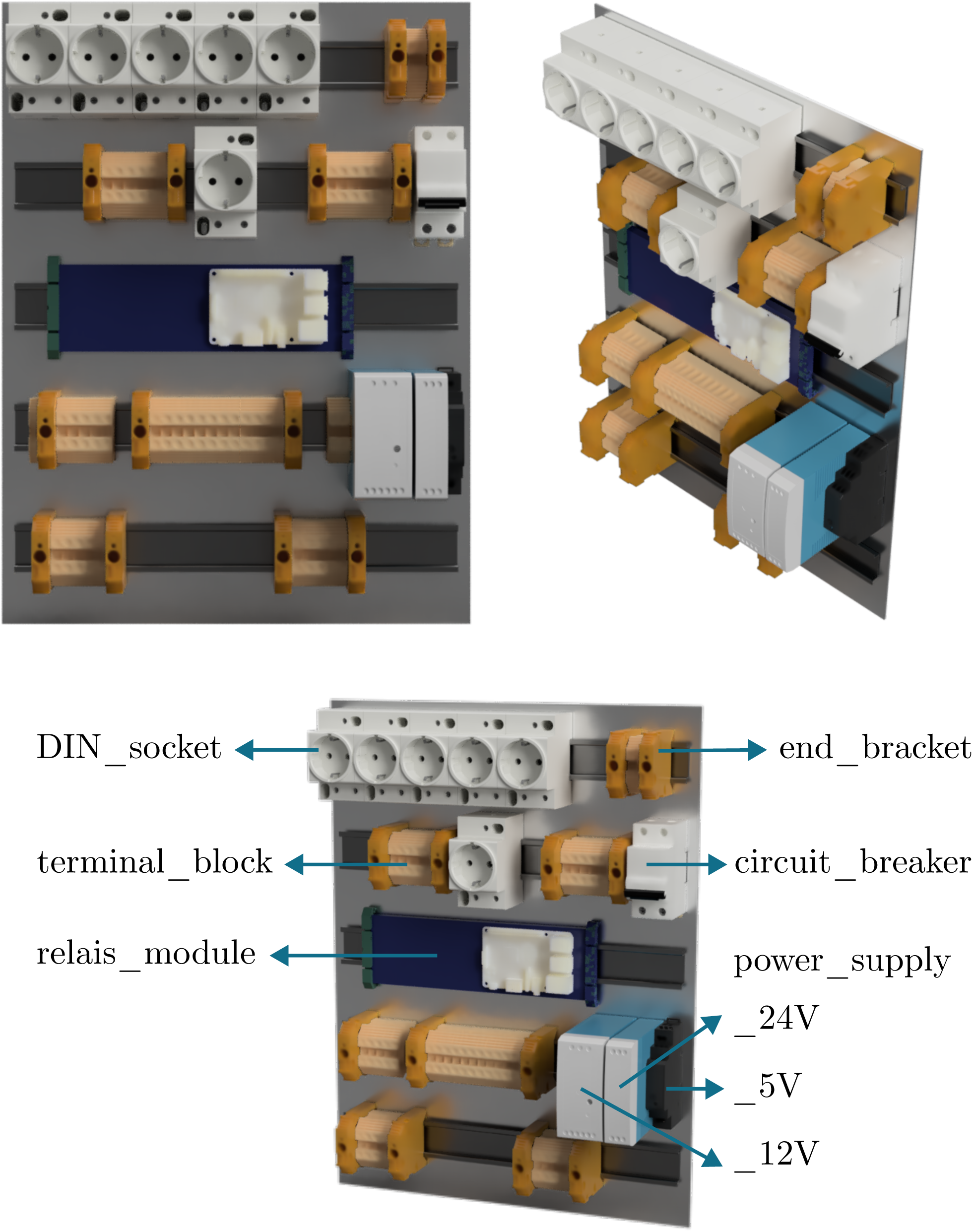
3D render of the proposed layout of the electrical cabinet as assembled in Figure 12. The annotations correspond to the part identifiers of the bill of materials.

### 5.4 Electrical cabinet (low-voltage)

The low-voltage circuitry (5 V, 12 V and 24 V) mainly consists of the Raspberry Pi and darlington arrays (2x `darlington_array`) to control the power to the thermoelectric condensers and the solenoid valves, as well as reading out the temperature probes. While you could also use relays for the low-voltage control actions, the darlington arrays are faster, cheaper, and more space-efficient. In general, all connections were made using rainbow wires (`rainbow_jumper_cables`) to keep the wiring organised and easily traceable.

1. Solder the darlington arrays, 4.7 kΩ resistor (`resistor`) and rectifier diode (4x `diode`, one for each valve) to a circuit board (`soldering_board`). An example layout of the board is shown in Figure 15.
2. Connect all components according to the design file `schematic_control`.

a. Ensure the GND of the (low-voltage) power supplies are connected.
b. The DS18B20 temperature probes all have a unique identifier which allows them to be connected in parallel to a single port of the RPi. Their only requirement is to connect the dataline to the 5 V line over the resistor, as indicated in the diagram. It is useful to use a 3-wire conductor (`wire_3_ conductors`) coming from the cabinet to limit cable clutter at the sensor side (using a 3-way terminal strip to connect several probes in parallel: `terminal_strip_3_way`).
c. Connect the first darlington array to the 5 V power supply and GND (similar to the relay board). The second array’s COM port should be connected to the 24 V power supply. In this way, they control the Peltier power supplies and the solenoid valves, respectively.
d. The control connections for the condensers are made to the remote ON/OFF (CN1) ports of the Meanwell power supplies. In this way, if the corresponding GPIO pin is set to high, the power supply will shut down. The 12 V output power is connected to the Peltier elements and the fans, in this case using 6 conductor wires (`wire_6_conductors`) for easier connection at the condenser side (using 6-way terminal strips, `terminal_strip_6_way`, see supplementary Figure S2). In principle, you could also solder them in parallel at the condenser side and apply only a two-wire conductor, but this is not ideal if you would need to service one of the components.
e. For the solenoid valves, extra rectifier diodes were installed to accommodate the inductive load. The diodes are connected with the cathode to the (24 V) COM port of the darlington array, and the anode to GND. In principle, one diode should be enough, but here one for each valve is used for safety. After all, voltage spikes can drastically shorten the lifespan or even destroy the electronics. In an initial design iteration, the gas valves were controlled with the relais board, but the integrated flyback diodes of the board were not sufficient to protect the RPi from voltage spikes, leading to short power interruptions every time the valves were switched off. The proposed diodes should therefore not be considered optional.
f. The exact connections to the GPIO pins of the Raspberry Pi are flexible, but should be considered alongside the limitations of the relay board. You can also cut the board, mount the RPi separately to the DIN rail, and then directly connect to the relais control terminals if you would need more pin flexibility.

### 5.5 Gas supply

The required gas supply to the system heavily depends on the application and laboratory environment. In the following, it is assumed that access to a central gas distribution system is available. If this is not the case, individual gas bottles need to be installed and replaced regularly. Be aware of the dangers of working with compressed gases, and always follow the safety guidelines of your institution. For example, in the case of nitrogen, asphyxiation is a relevant risk, and ensuring the ventilation system is functioning properly is of vital importance. The following steps are a general guideline to set up the gas supply for the BIO-SPEC system.

1. For every type of gas, add a pressure regulator (`pressure_regulator_main`) to the gas line and adjust to 2 barg to limit the maximum pressure in the system and avoid fluctuations. Apply 6 mm PTFE tubing and push-in fittings (`push_in_main`) to make the connection.
2. In case of a centralised compressed air supply, a filter (`pneumatic_filter`, with 2x `push_in_individual`) should be added after the regulator to remove any oil droplets coming from the compressor.
3. Connect the gas supply to the solenoid valves (4x `solenoid_valve_gas`) using PTFE tubing. In the proposed design, two sets of reactors can be separately sparged with air or nitrogen, which is why four solenoid valves are connected as shown in Figure 16a, using four Y fittings (4x `push_in_Y`).

a. Note that theoretically, in this configuration, you could push one type of gas into the other gas line if you would accidentally open all solenoid valves at the same time. If your system does not have any check valves to prevent this, it is recommended to add them yourself (2x `check_valve_gas_supply`) after the main pressure regulator. Most regulators have a similar feature built-in, but may not guarantee a perfect seal at the incidental backward pressure. Any accidental gas contamination is not worth the minor investment in these check valves.
b. The solenoid valves only open when power is supplied. As such, they are normally closed (NC), which means that in case of a power failure, the gas supply is automatically shut off. Hence, the loss of power to the thermoelectric condensers is not a safety hazard in this situation.
4. Every reactor then needs a separate pressure regulator (`pressure_regulator_individual`, with 2x `push_ in_individual`) and rotameter (`rotameter`, with 2x `push_in_rotameter`) to keep the gas flow to each vessel as consistent as possible.

- The rotameter should be set to the desired flow rate (at least 1 vvm is recommended), and the pressure regulator adjusted to 300 mbarg (using `pressure_gauge`), to avoid shattering the reactor vessel if a pressure build-up would occur during operation. If the pressure is not controlled individually for each rotameter, the gas flow will be distributed according to the resistance of the tubing, reactor, filter etc., which can lead to a varying gas injection across replicates.
5. To easily stack the pneumatic components, an optional box was designed (design files `plate_regulators_ rotameters_5mm_I` and `plate_regulators_rotameters_5mm_II`) that can be lasercut in 5 mm plexi. The assembly is shown in Figure 16b.
6. Add an extra manual valve (`manual_gas_valve`) after the rotameters, to be able to shut off the gas supply to the reactor vessel without having to adjust the rotameter every time. This valve is especially useful when the reactor vessel needs to be disconnected from the gas supply. The valves can also be fixed in place using an optional lasercut plexi box (`plate_valves_3mm`) as shown in Figure 16c.

**Figure 15:**
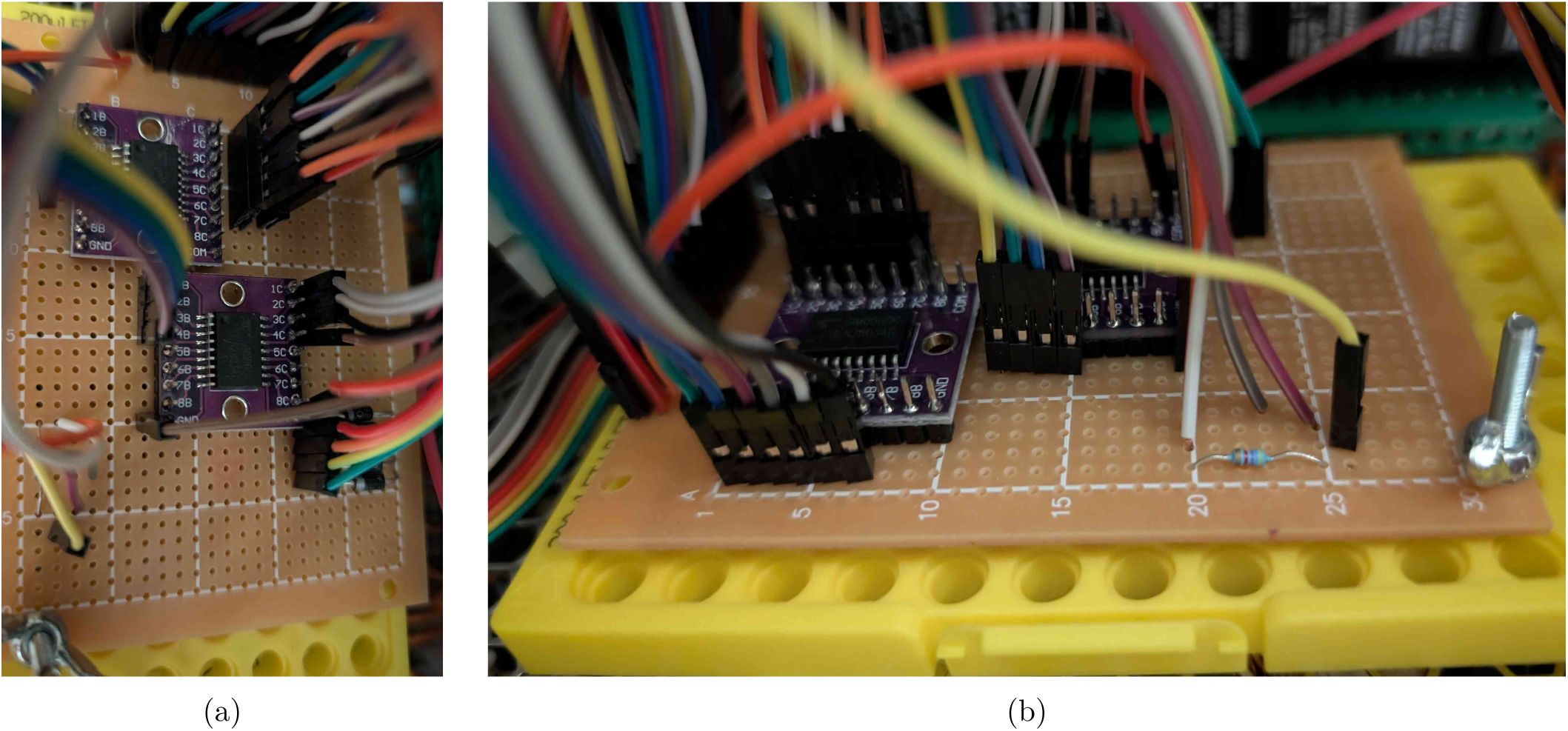
Top and side view of an example layout of the circuit board for the low-voltage electronics.

**Figure 16:**
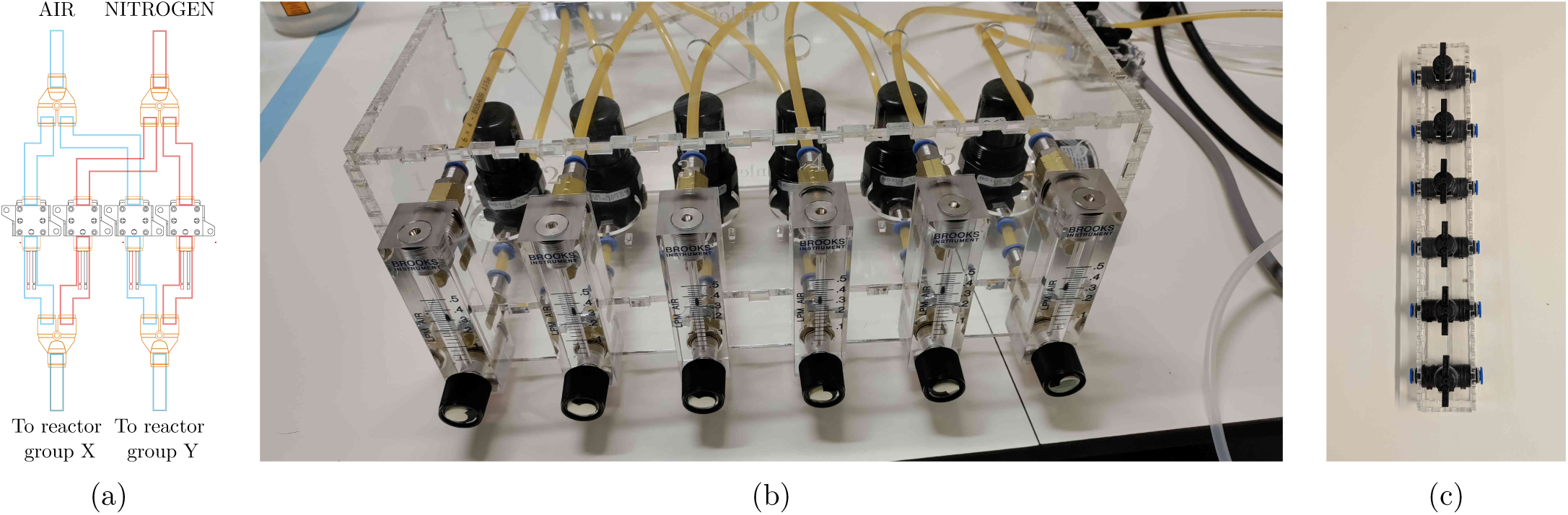
Pictures and diagrams of how to set up the gas supply for two sets of three reactors. (a) shows the solenoid valve connections, (b) the rotameter and pressure regulator box, and (c) the plexi box for the gas valve array.

### 5.6 Setting up the Raspberry Pi

As a last step before operating the BIO-SPEC system, the control software (`main_control.py`) needs to be loaded on the Raspberry Pi (`rpi_board`). Evidently, this is the most flexible part of the system, and you can adapt the software to your needs. An Arduino could be useful as well, but having an HMDI monitor as well as a keyboard and mouse is much more convenient for the intended type of experiments. In addition, by using Python, a simple GUI can be created to more easily monitor what is going on during operation. An alternative control script with continuously updating graphs is provided as an extra example of how you can extend the software (`main_control_graph_example`). For the majority of applications, the standard script is expected to be adequate.

1. Download the latest Raspberry Pi OS image from the Raspberry Pi website and flash the image to an SD card (`rpi_SD_card`) using a tool like Etcher.

a. Enable the ‘One-Wire’ interface to read out the DS18B20 temperature sensors by adding the following line to the `/boot/config.txt` file and rebooting the RPi: to ve r l a y=w1−g pio
b. Enter the following commands in the terminal to retrieve the unique identifiers of any connected temperature probes: $ sudo modprobe w1−g pio
$ sudo modprobe w1−therm
$ **`cd`** / sys / bus/w1/ d e v i c e s
$ l s
The unique identifiers will be listed here, one directory for each probe. Connect them one by one, writing down the identifiers of the form “28-X*”. You will need to change these in the code later on to match your unique probe for each reactor. Note that the sensors are connected in parallel and read out in bulk to reduce readout time. In addition, the temperature resolution is reduced to 11 bit at the start of the script to get the minimum required accuracy of the reading at a higher refresh rate. You can already test the resolution and bulk readout by entering the following command in the terminal (changing 28* with the relevant identifier):
$ **`echo`** 11 *>* / sys / bus/w1/ d e v i c e s /28∗/ r e s o l u t i o n
$ **`echo`** t r i g g e r *>* / sys / bus/w1/ d e v i c e s / w 1 bus master 1 / therm bulk read
$ **`cat`** / sys / bus/w1/ d e v i c e s /28∗/ temperature
c. Install the necessary Python packages using requirements.txt. This can be done by connecting the RPi to the internet and running: $ pip i n s t a l l −r requirements .txt
  i. You can of course use your favourite virtual environment manager to keep the Python packages separate from the globally installed packages. Do keep in mind that you will then need to change the path to the Python interpreter and/or activate the environment in the startup script (see further).
d. The control script can be started by running: $ python3 / path / to / mainloop T read .py
e. As an extra safety feature, the control script should be started automatically after booting the Raspberry Pi. This can for example be achieved using (root’s) crontab to call a bash script `startup_ script.sh` after boot. $ sudo crontab −e
Add the following line to the crontab file.
@reboot / bin / s l e e p 2 0; DISPLAY=:0 bash path / to / s t a r t u p s c r i p t .sh
The bash script should then contain the code below. First, the necessary kernel modules for the temperature sensors are loaded, and the Python control script is started. Any additional output is redirected to a log file, which can be useful for debugging.
*#! / usr / bin / bash*
modprobe w1−g pio
modprobe w1−therm
/ usr / bin / python3 / path / to / main control .py *>>* / path / to / l o g .txt
In this way, if power is lost, the system will automatically restart the script after booting and continue the experiment. For this reason, any starting conditions should be hard-coded. For example, to determine the current reactor conditions that need to be applied, the Python code should under no circumstances rely on the time the script was started.

i. You may also consider calling the startup script from `/etc/rc.local` instead of using crontab. The script is then also run as root after boot, and should therefore also not give you any issues with permissions.
f. After setup, you do not need to connect the RPi to the internet any more, as the script will run independently. If you do wish to keep the hardware connected, you should take care to secure it properly.

## 6. Operation instructions

### 6.1 Setting up the software

The `main_control.py` script is designed to be as flexible and intuitive as possible, since it will require some changes to match your specific setup and experimental design. It is important to test your program before every experiment, as discussed below. In addition, when adapting to software to the user requirements, the `tutorial_main_control.html` file provides an extended explanation of the code and implementation decisions to improve clarity.

1. Ensure that the correct GPIO pins are defined in the script, and that the unique identifiers of the tem-perature probes (`device_keys_bulk`) match the corresponding power sources of the Peltier elements (in the `temperature_PROBE_PIN` list).
2. In the provided code, up to six reactors can be controlled (index 0-5). The ones that are not in use can be given a random probe key, since the script will only read out the temperature probes that are connected to the RPi anyway. Every probe that is not found will just give a constant temperature reading of *−*5 °C to avoid turning on unused power supplies. For 6 reactors, with only 4 connected, the setup would look like the lines below. temperature PROBE PIN = [ IO pin (pinnumber=12, pintype=” p e l t i e r “), *# R1*
IO pin (pinnumber=10, pintype=” p e l t i e r “), *# R2*
IO pin (pinnumber=8, pintype=” p e l t i e r “), *# R3*
IO pin (pinnumber=13, pintype=” p e l t i e r “), *# R4*
IO pin (pinnumber=15, pintype=” p e l t i e r “), *# R5* IO pin (pinnumber=11, pintype=” p e l t i e r “), *# R6*

]
d e v i c e k e y s b u l k = { ”/ sys / bus/w1/ d e v i c e s /28 −00000 ec 61 a 33 / temperature “ : 0,
”/ sys / bus/w1/ d e v i c e s /28 −00000 ec 60 edb / temperature “ : 1,
”/ sys / bus/w1/ d e v i c e s /28 −00000 e 66 bec 8 / temperature “ : 2,
”/ sys / bus/w1/ d e v i c e s /28 −00000 ec 7 b 190 / temperature “ : 3,
” 28 x” : 4,
” 28 y” : 5,
}
3. The directory `RPi_debug` contains a python file `GPIO.py` that can be used to test the intended cycles and control actions on a computer before running it on the Raspberry Pi. Add the directory to the same working directory as `main_control.py` and simply comment the line: import RPi. GPIO as GPIO while uncommenting the import of the test module. import RPi debug. GPIO as GPIO
4. Define the desired timing (’interval’, as a ‘Pandas Timedelta’ object) for the phases that you want to apply using the ‘schedule’ class, and also specify the ‘START TIME’ of the experiment. The ‘phase’ module also defines in which state the sockets (for stirrers and pumps) and solenoids need to be in for every individual phase. These definitions need to be changed to match your needs. An example for a stirred phase with continuous nitrogen sparging is given below.

command dict = { ”AIR” : False,
”N2” : True,
”FEED” : False, “FEED2” : False,
”OUT” : False,
”STIR” : True,
}
(a) Note that two sets of reactors, referred to by their ‘groupkey’ (’X’ and ‘Y’), can be controlled separately. This is useful if you want to compare two different cycling conditions, or if you want to run two different experiments at the same time. Of course, this separation can be extended to more groups if needed.
(b) The ‘schedule’ class is also implemented with an initial ‘growth’ phase (defined by ‘TIMESTAMP GROWTH’), where (in this case) an aerobic mixing regime is operated until a predefined point in time before cycling ensues. If you would not need this, simply set it to match the start time of the experiment.
(c) It is useful to uncomment the print statements in the ‘phase’ module to check the phase transitions and timings beforehand. This check is easily achieved by setting the phase intervals to seconds and checking if everything loops as expected, and the right equipment is turned on/off at the right time. Make sure to comment the print statements again before the actual experiment, since this drastically slows down the control script. The complete cycle is also visualised automatically in the GUI by the changing color and text of the buttons, as illustrated in Figure 17.

**Figure 17:**
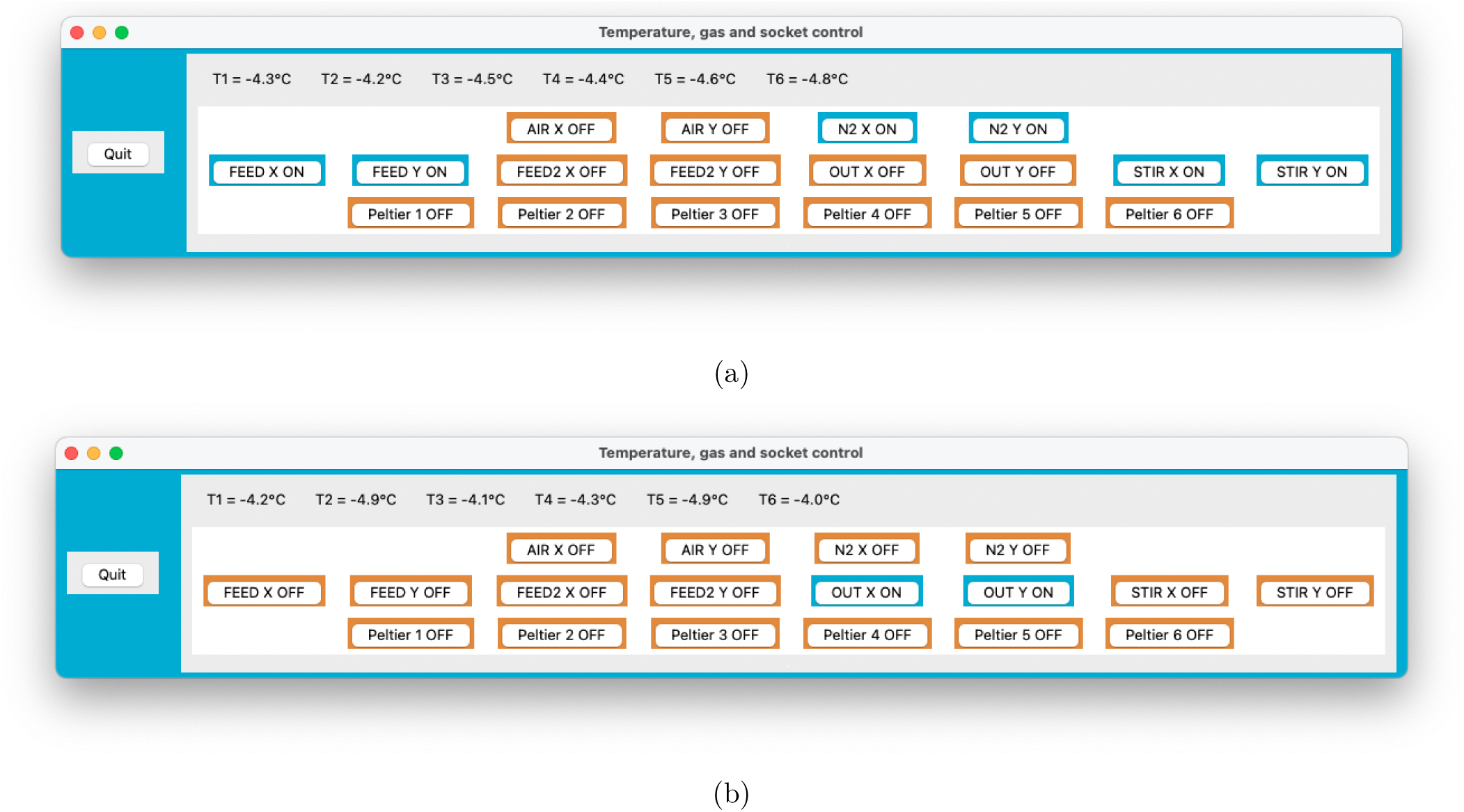
Screenshots of the GUI during an anaerobic feeding (a) and a withdrawal (b) phase. The buttons change their color (BLUE/ORANGE) and text (ON/OFF) according to the current phase. The Peltier elements are all turned off here, since this was run without any probes connected.

### 6.2 Preparing the vessel

1. Autoclave the reactor vessel, medium, and all tubing connections.

a. The silicone tubing (e.g., `silicone_tubing_3×6` and `silicone_tubing_4×7`) should be cut to length beforehand and should contain parts with peristaltic pump tubing as well (connected with luer locks). Here, only the parts directly connected to the vessel and bottles are of size 4 mm *×* 7 mm (IDxOD) to prevent nicks in the tubes, while the rest is 3 mm *×* 6 mm to limit the dead volume.
b. The headplate can be autoclaved separately (covered in aluminium foil) if the complete vessel would be too large for your autoclave.
c. Keep in mind that just like the reactors, any feed or waste bottle also needs to be fitted with a sterile filter. These filters do not only avoid contamination, but also even out any pressure build-up in the system due to pumping (as long as they are not wetted). If you do not have any two- or three-port headplates for general GL45 bottles, the design files `GL45_threeport_UNF` and `GL45_twoport_UNF` allow you to manufacture your own (compatible with `UNF_barb_fitting_Masterflex` and `O_ring_ GL45`).
d. If your medium precipitates during autoclaving, it is recommended to either combine the components separately after autoclaving, or sterilise them by filtration to prevent clogging.
e. For feeding bottles, it is recommended to make a cut-out at the bottom of the inner tube, to prevent the tube from getting stuck against the bottle wall (due to suction) and blocking the flow.
2. After sterilisation, add the medium to the reactors and make the necessary luer-lock connections near a flame or in a laminar flow cabinet, as shown in Figure 18.
3. Inoculate the reactors either by opening the lid, or preferably through a septum or sample valve. As shown in Figure 19a, an autoclavable blunt needle is suitable to inject the culture through the valve without having to open the reactor.

a. This step could also be performed outside the biosafety cabinet if needed.
b. After inoculation, you can reuse the valve to for example spike the content manually with medium/acid/base/… by applying a standard 0.2 µm filter (see Figure 19b). If you manually add acid and base through the same port, dilute them first (ideally below 1 M), and flush the port afterwards with water to avoid the hazard of mixing concentrated acid and base. In any case, the dilution and flushing steps are recommended to limit corrosion.
4. Place the vessels on magnetic stirrers and connect the tubing to the peristaltic pumps.
5. Prime the tubing (remove all air) by filling the feed lines to each reactor. If all lines are of equal length, you can prime them at the same time.
6. Connect the water bath to the reactor vessels using GL18 hose barb connectors (included with the vessel) and silicone tubing (e.g., `silicone_tubing_8×12`), with the inlet at the bottom, and outflow at the top. Ideally, the reactors should be connected in parallel (with tubing of equal length) to avoid any temperature gradients. Though during experiments at 30 °C, two pairs of two reactors in series did not differ in temperature more than 1 %, which was fine for our application.
7. Connect the thermoelectric condensers and temperature probes to the reactors.
8. Connect the gas inlet of each reactor to its sterile filter using PTFE tubing, and either silicone tubing at the end (to fit over the hose-barb), or a push-in reducer (`push_in_reducer`).
9. Plug in the electrical cabinet to start the control system, the power to the condensers and to activate the gassing system.

### 6.3 Shutting down the experiment

1. Quit the control script by pressing the ‘Quit’ button in the GUI, or by pressing ‘Ctrl+C’ in the terminal.
2. Shutdown the RPi using the command: $ sudo shutdown now
3. Disconnect the power to the electrical cabinet and detach the thermoelectric condensers. Alternatively, you can just press the emergency button.
4. Close the manual gas valves and disconnect the gas supply to the reactors.
5. Disconnect the water bath from the reactors. Ideally, you should put the water bath on a lower height, and slowly unscrew the upper GL18 port to allow air to enter the circuit and drain the reactors one by one without spilling. In this way, you can also reuse the cooling water. If you notice algae growth in the water bath, you may need to add some algae inhibitor to the water (available at any aquarium store). Do not use bleach, as this will corrode the pump of the water bath.
6. The reactors (and waste bottles) should be autoclaved, disposed of the broth, and cleaned thoroughly as follows.

- Disconnect the tubing from the reactors and rinse with water. You can reuse the parts of the tubing if no biofilm is present, but it is recommended to replace it after a few experiments.
- Soak all glass, stainless steel and polypropylene components in alkaline soap (e.g. RBS^TM^ 35, `RBS_ soap`). Under no circumstances should you put the aluminium off-gas pipe in the same bath, as this will corrode extremely quickly in alkaline solutions.
- Clean all components with distilled water and a brush (a toothbrush works really well), and make sure no soap is left after rinsing.
- Double-check the integrity and cleanliness of the O-rings and replace if necessary.
- Dry, assemble, and autoclave the reactors again for the next experiment.

### 6.4 Troubleshooting

To guide the user during long-term operation, some common issues as well as relevant troubleshooting experience are summarised in Table 3.

**Figure 18:**
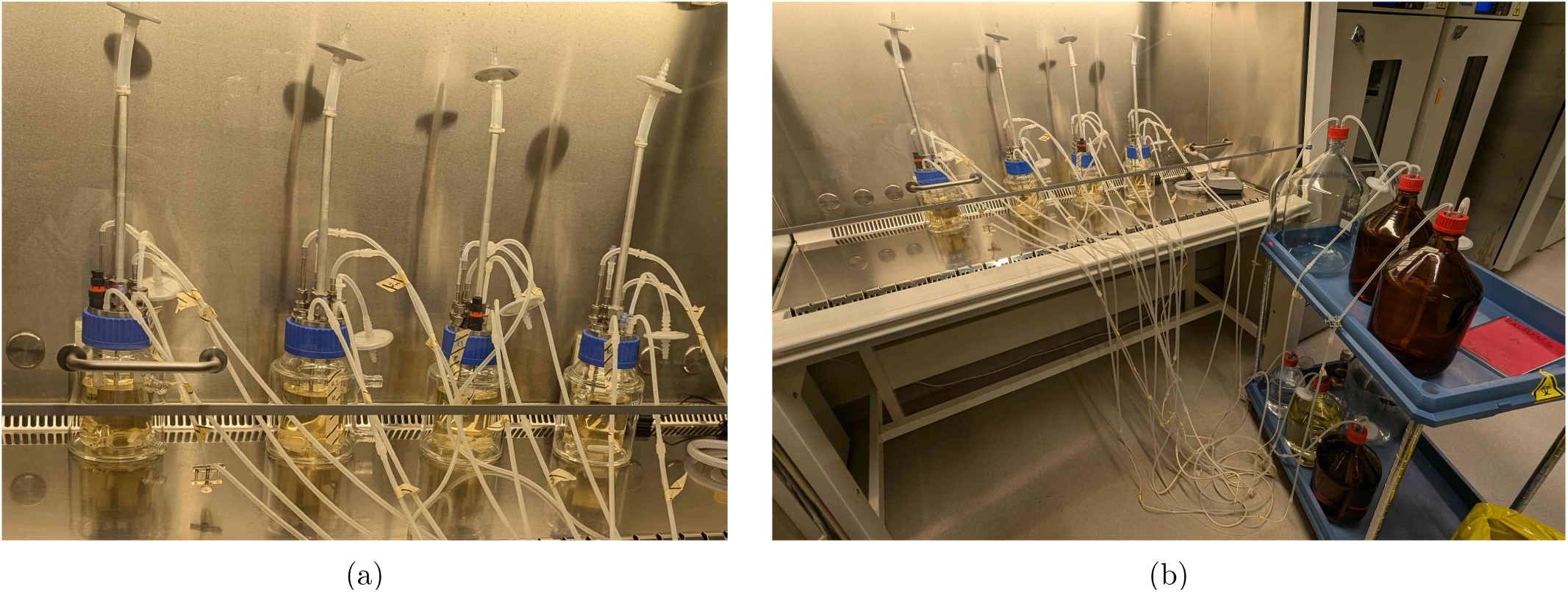
Reactors are connected to feed and waste bottles using luer locks in a laminar flow cabinet. In the setup shown here, every reactor is connected to an individual 5 L feed bottle, and the waste lines are interconnected after the peristaltic pump tubing to two 10 L waste bottles in series.

**Figure 19:**
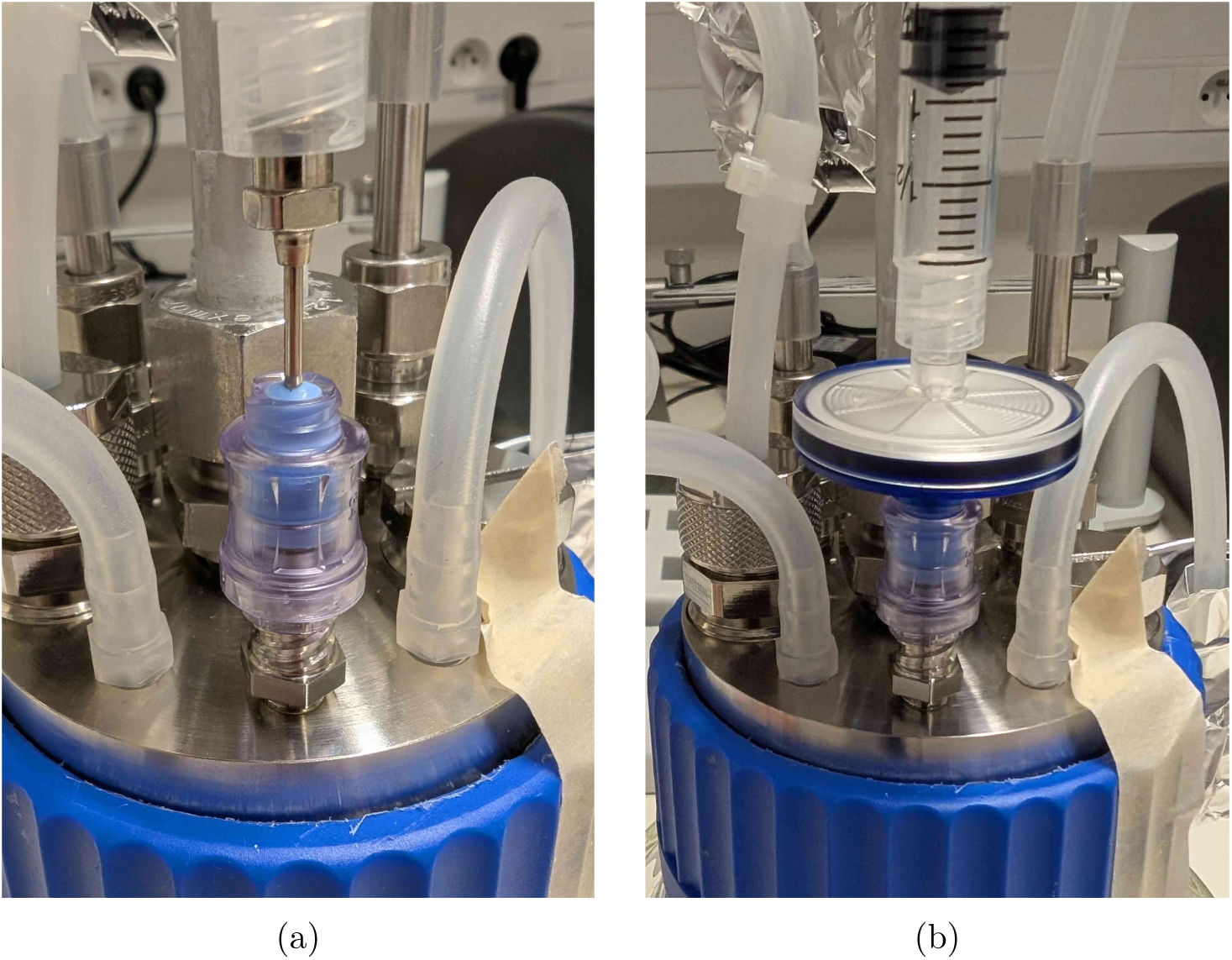
Inoculation and feeding options for the sampling valve. The blunt needle applied in (a) keeps the valve intact to allow the use of the same port for the manual addition of spiking solutions through a 0.2 µm filter in (b).

**Table 3:** Common issues and their solution strategy during operation of the BIO-SPEC system.

| Problem and possible causes | Suggested course of action |
| --- | --- |
| Temperature reading of the condenser is not around the setpoint.<br>• Not enough cooling power due to a faulty Peltier element.<br>• Power supply overheating.<br>• Control application froze. | Shut down the gas supply by closing the rotameter.<br>• Check the connections of the 6-way terminal strip.<br>• Make sure the temperatures are still updated in the GUI. If they are not, press quit and restart the control script.<br>• Check the LED indicator of the corresponding power supply. If it is bright green, there is no issue. However, if it is faint, the system is not receiving the full power capacity due to overheating of the power supply. Install an extra cooling fan and check if that improves the situation. If the issue persists, consider replacing the power supply.<br>• Detach the condenser unit and remove the fans from the assembly. Unscrew the aluminium plate to access the Peltier elements. Connect a 12 V battery to the Peltier elements, ensure their temperature drops at the condenser side, and replace the unit if it does not cool down any more. Always inspect faulty thermoelectric coolers for cracks, since overtightening the assembly may have been the root cause of that. During reassembly, make sure the thermal pads are intact. |
| Reactor leaking or losing air after assembly. | Tighten fittings securely. Before autoclaving, inspect O-rings for damage or wear and replace if necessary. During assembly, ensure that they are compressed uniformly. |
| Temperature probes not detected by the Raspberry Pi | Verify the unique identifiers of the probes and check the wiring connections. |
| Reduced gas flow at the same inlet pressure.<br>• Off-gas filter clogged over time.<br>• Off-gas filter clogged due to foam formation.<br>• Biofilm formation on the aerator. | • Replace the off-gas filter or install parallel filters to more easily switch between them. In case of the former, it is recommended to work in a biosafety cabinet or to first increase the gas flow rate to the maximum to reduce the risk of contamination.<br>• If your medium produces a lot of foam when aerated, consider adding an antifoam agent (e.g. Antifoam 204) at regular intervals.<br>• If a lot of biofilm is formed over time, consider increasing the gas flow or stirring rate if possible. In between runs, make sure to flush the aerators thoroughly with water, soak in isopropanol, and do not make skin contact with the aerator surface to avoid the accumulation of lipids. |
| High memory usage of the Raspberry Pi after multiple weeks of operation. | Check the accumulated memory of the 'lxpanel' process and kill it to free up space. This process is responsible for the desktop panel and will restart automatically. |

## 7. Validation and characterisation

The BIO-SPEC was validated in a series of experiments with co-cultures of bacteria. Before diving into the results, the main capabilities and limitations of the system are summarised below.

- A wide range of applications can make use of the proposed setup, from processes requiring simple batch experiments to complex cycling regimes in terms of gas and liquid flow. All aspects of the system are modular and customisable for this exact reason.
- Consistent experimental conditions can be applied to (up to) six replicate reactors, or for example two sets of (up to) three reactors. The feed, gas supply, and stirring can be separately controlled for each set, which means two conditions can be compared in parallel, or two experiments can be run at the same time. If needed, it is also possible to extend the number of reactors or sets, requiring extra hardware or software respectively.
- The system is able to control the temperature of the thermoelectric condensers and all other peripherals for at least 35 days without downtime. During operation, the software allows for real-time monitoring of the applied reactor phase through a GUI. In addition, it ensures safe operation even in case of a power interruption.

**–** After a few weeks, fats, oils and volatile components from the broth accumulate on the off-gas filter, leading to a reduced gas flow. Therefore, for long-term operation, more parallel filters should be installed on each pipe, or the filter should be replaced regularly under sterile conditions (which implies downtime of a few hours).
**–** The estimated cost of the system is relatively low, but it does not include the cost of the pumps, sensors, and other peripherals, since these are highly dependent on the requirements of the intended application. Overall, the glassware and fittings are the most expensive components of each vessel, but they can be reused indefinitely.
**–** The accuracy of the hydraulic retention time is limited by the available peristaltic pumps. In addition, with simple on/off control of the pumps, it is not possible to vary the flow rate during operation, which might be a limitation for some applications. As a workaround, several feeding lines with different tubing dimensions can be used in parallel to achieve a wider set of flow rates, but switching those lines cannot be done automatically without additional hardware. The previously mentioned FAST pump project [11] could provide an alternative way out though, since the stepper motor is directly controlled by the RPi and could therefore be programmed to change the flow rate during operation.

To validate the system, two coculture experiments with bacterial strains that are relevant in a biological wastewater treatment context were set up. On the one hand, *Azoarcus communis* Swub 3 (DSM 12120), *Tetrasphaera japonica* (DSM 13192), and *Sphaerotilus montanus* (DSM 17940), were used in sequencing batch reactor (SBR) operation. The applied cycles consist of an anoxic feeding, anoxic denitrification (stirred), aerobic (stirred), settling, and withdrawal phase. The base medium and metabolite analyses are described by Parret et al. [15], while nitrate and glucose were used in the feed instead of ammonium and acetate. The most relevant reactions here are the fermentation of glucose to acetate (Equation 1), the aerobic respiration of acetate (Equation 2), and the two-step denitrification from nitrate to nitrogen gas, over nitrite (Equation 3).

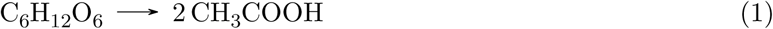

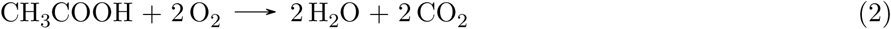

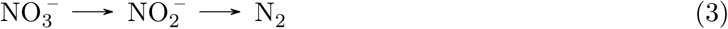

To facilitate all reactions, both oxic and anoxic conditions need to be applied by sparging air and nitrogen, respectively. The results for two cycles are shown in Figure 20, and clearly show the feeding and gas control of the BIO-SPEC.

**Figure 20:**
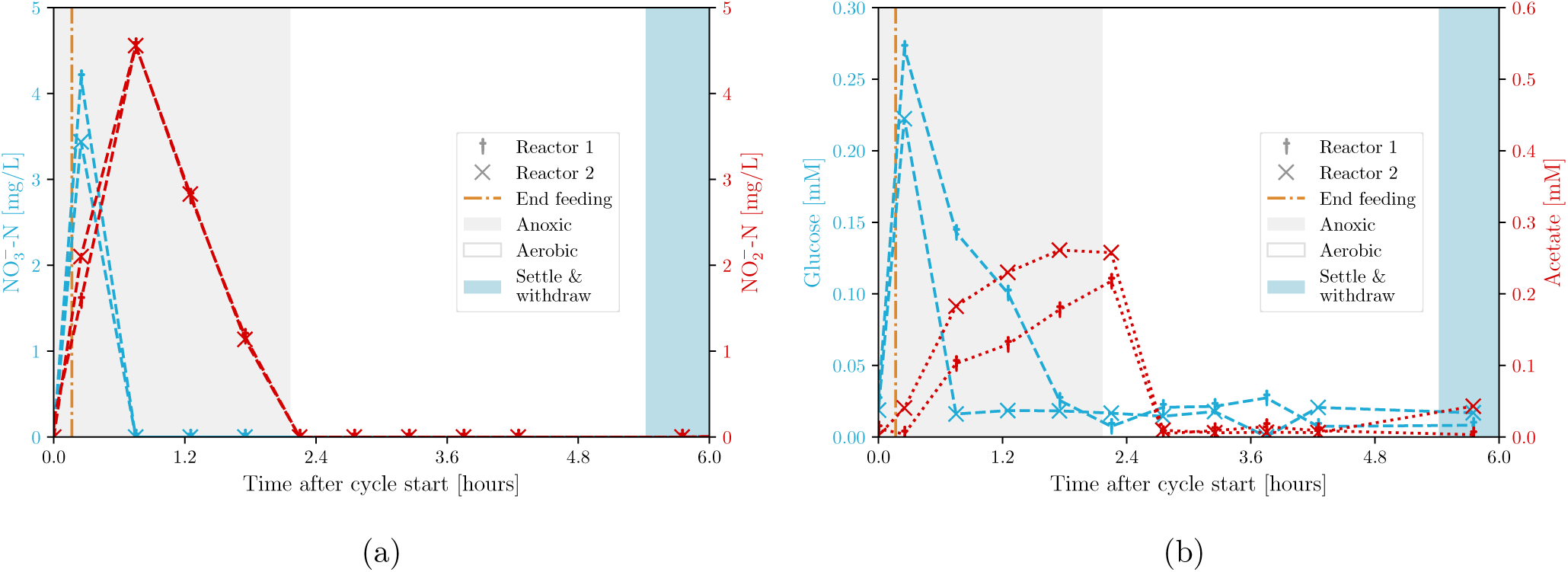
Metabolite concentrations produced during anoxic/oxic cycles in a coculture experiment for two replicates. Denitrification (Equation 3), fermentation (Equation 1), and respiration (Equation 2) are observed. Notice the accumulation and removal of nitrite as an intermediary product during the anoxic phase, and the immediate consumption of acetate in aerobic conditions.

On the other hand, during a five-species experiment (now also including the nitrifiers *Nitrosomonas europaea* (DSM 28437) and *Nitrobacter winogradskyi* (DSM 10237)), the complexity was extended to include the two-step nitrification reaction as well (Equations 4 and 5).

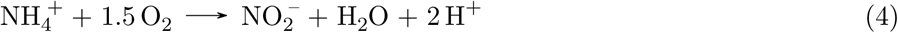

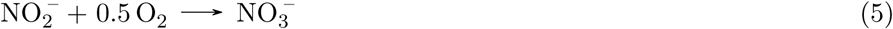

A second feed with extra carbon source was also added after the aerobic phase to facilitate denitrification (and not respiration). The results shown in Figure 21 indicate the progressive loss of nitrifier biomass (and therefore nitrification) over several days. Since nitrifiers grow slowly and do not settle compared to the other organisms, these results do not only demonstrate the correct control of feeding and gas supply again, but also of the previously implied withdrawal phase. After all, planktonic biomass is just transported to the waste vessel during withdrawal, lowering their abundance. In conclusion, the BIO-SPEC can be used to study the dynamics of complex microbial communities in a controlled environment during long-term cultivation.

**Figure 21:**
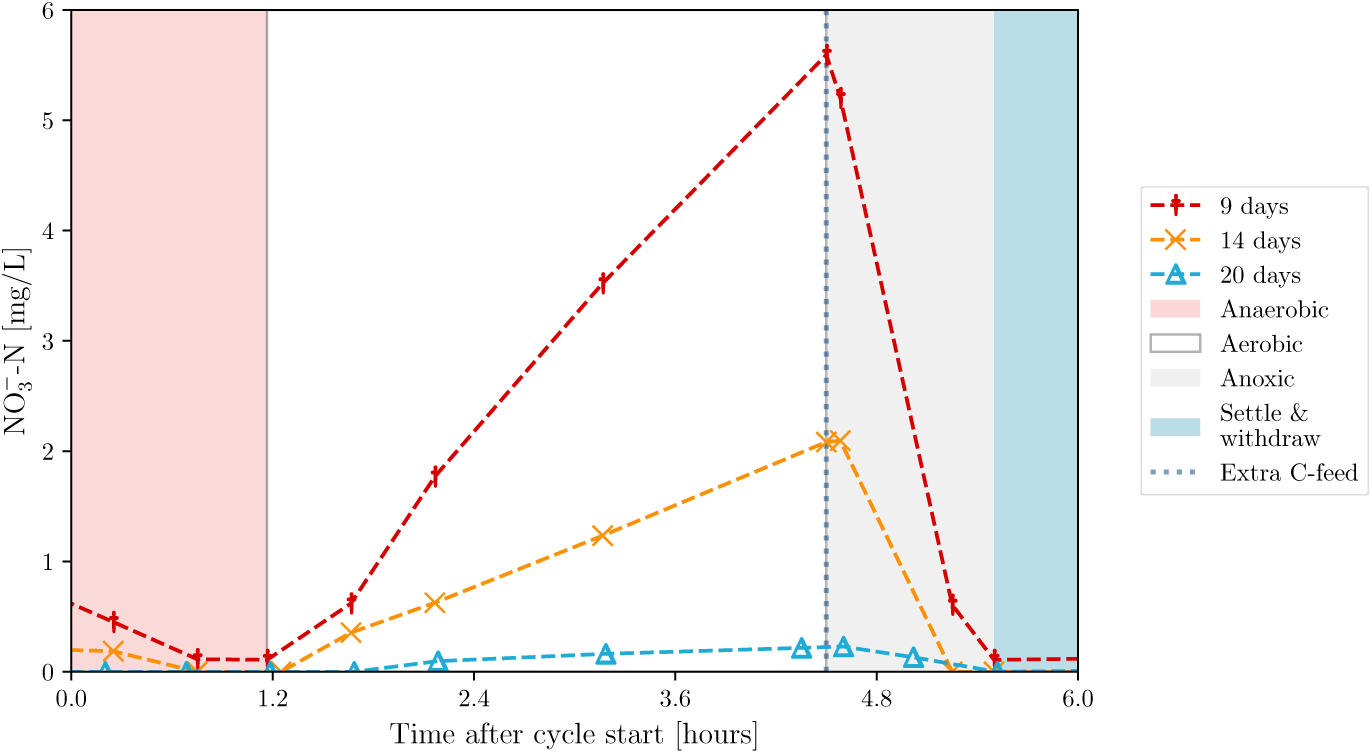
Nitrate profiles for one reactor of a coculture experiment in SBR operation. A similar decline was observed in all four replicates, indicating the loss of nitrifiers over time.

### 7.1 Future improvements and extensions

The BIO-SPEC is a modular system that can be adapted to a wide range of applications. The improvements or extensions that are relevant to further enhance its capabilities are summarised below.

- **Integrate open-source peristaltic pumps (FAST pump project** [11]). Enable precise control of feeding rates, sampling, or waste removal. This integration would also allow for dynamic flow rate adjustments during operation.
- **Include pH control.** Add pH probes with integrated chips to more easily read and send the pH signal to the RPi controller. For non-sterile setups, cheaper pH probes that are directly compatible with the RPi can be used. Either way, the software needs to be expanded to add acid or base to the reactor, similar to how temperature control is currently implemented.
- **Include mass flow control.** Include mass flow controllers for precise gas flow regulation when a more delicate application is envisioned. Bronkhorst mass flow controllers can be integrated using the “Bronkhorst Propar” Python package.
- **Design a more compact condenser.** To reduce footprint, a smaller heat sink could be used. However, this would also require a more powerful thermoelectric cooler (and power supply) to deliver a higher temperature difference between the two sides of the condenser. After all, the hot side will be warmer when applying a smaller heat sink surface area. If room temperature can be controlled, however, a smaller heat sink may still be adequate for safe operation. In general, if a chiller is already available, use the proposed coolant-based condenser to simplify the circuitry and eliminate the need for active temperature control.
- **Use a larger electrical cabinet.** Enhance the cooling of the electrical cabinet by adding more fans and using a larger cabinet to prevent overheating of power supplies. In a larger cabinet, there is also more space to fit the components and more options to expand the hardware.
- **Install parallel off-gas filters.** Reduce downtime and contamination risks caused by required maintenance of clogged air filters during long-term experiments with foaming media.
- **Extend reactor sets.** Scale the system to accommodate more reactors or additional experimental conditions by expanding the hardware and reactor sets in the software.
- **Improve manual interactions with the hardware.** Extend the code to allow for easier manual interventions to change the operating conditions during the experiment. Right now, the script needs to be restarted if certain cycle parameters need to be adapted. This could be improved by adding a manual control mode, where the user can change these predefined parameters in the GUI.
- **Add scales to the reactors.** A lot of lab scales are equipped with serial RS-232 connections (e.g. A&D EK-i series). This means that the scales can be connected to the RPi using a RS-232 to USB converter, and the data can then be read using the pySerial library. As such, real-time monitoring of the reactor mass is possible, which is useful to control the feed and/or waste pumps. Not only would this extension avoid the need to calibrate peristaltic pumps, it would also allow for more precise control of the hydraulic retention time.

## Conflict of Interest

The authors declare that they have no known competing interests.

## Ethical approval

This article does not contain any studies with human participants or animals performed by any of the authors.

## CRediT author statement

**LP**: Conceptualisation, Methodology, Investigation, Writing - Original draft preparation, Visualisation. **KS**: Conceptualisation, Methodology. **JDV**: Writing - Review & Editing, Supervision, Funding acquisition. **IS**: Writing - Review & Editing, Supervision, Funding acquisition.

## Acknowledgements

This work was supported by KU Leuven [C24/18/043] and the Research Foundation Flanders [FWO-G032321N]. Laurens Parret holds a FWO PhD fellowship Fundamental Research of the Research Foundation Flanders [FWO-1191022N]. The funding sources had no involvement in the study or its publication.

The authors would also like to thank Kristel Bernaerts for her support and help with safety features of the system, Jonas Van Olmen for his valued recommendations on the electrical hardware, and Wannes Van Holm for printing the headplate in autoclavable resin.

## Notes

https://doi.org/10.17605/OSF.IO/39WSB

### Competing Interest Statement

The authors have declared no competing interest.

## References

[1] A. Berenjian, Ed., Essentials in Fermentation Technology. Cham: Springer International Publishing, 2019. ISBN 9783030162306 Description based on publisher supplied metadata and other sources.

[2] W. Daniels, J. Bouvin, T. Busche, C. Rückert, K. Simoens, S. Karamanou, L. V. Mellaert, OÓ. H. Fridjónsson, B. Nicolai, A. Economou, J. Kalinowski, J. Anńe, and K. Bernaerts, “Transcriptomic and fluxomic changes in *Streptomyces lividans* producing heterologous protein,” Microb. Cell Fact., vol. 17, no. 1, dec 2018. doi: 10.1186/s12934-018-1040-6

[3] K. Turksen, Ed., Bioreactors in Stem Cell Biology: Methods and Protocols, 2nd ed., ser. Springer protocols. New York, NY: Springer US, 2022. ISBN 9781071620182

[4] N. Kapoor, E. Ndungo, L. Pill, G. Desalegn, A. Berges, E. V. Oaks, J. Fairman, and M. F. Pasetti, “Efficient production of immunologically active *Shigella* invasion plasmid antigens IpaB and IpaH using a cell-free expression system,” Applied Microbiology and Biotechnology, vol. 106, no. 1, pp. 401–414, Dec. 2021. doi: 10.1007/s00253-021-11701-4

[5] L. Bircher, A. M. Sourabíe, M. Paurevic, J. Hochuli, A. Geirnaert, C. Navas, B. Drogue, and C. Lacroix, “*Faecalibacterium duncaniae* A2-165 growth is strongly promoted by yeast extract and vitamin B5 in cGMP medium,” Microbial Biotechnology, vol. 17, no. 1, Nov. 2023. doi: 10.1111/1751-7915.14374

[6] T. Wenzel, “Open hardware: From diy trend to global transformation in access to laboratory equipment,” PLOS Biol., vol. 21, no. 1, p. e3001931, Jan. 2023. doi: 10.1371/journal.pbio.3001931

[7] B. G. Wong, C. P. Mancuso, S. Kiriakov, C. J. Bashor, and A. S. Khalil, “Precise, automated control of conditions for high-throughput growth of yeast and bacteria with eVOLVER,” Nat. Biotechnol., vol. 36, no. 7, pp. 614–623, Jun. 2018. doi: 10.1038/nbt.4151

[8] A. Graham, C. Thompson, D. Flynn, H. Elchos, J. Gibson, L. B. Priddy, and M. W. Priddy, “Design and construction of a low-cost compressive loading and perfusion flow bioreactor,” HardwareX, vol. 19, p. e00565, Sep. 2024. doi: 10.1016/j.ohx.2024.e00565

[9] J. Meneses, S. R. Fernandes, J. C. Silva, F. C. Ferreira, N. Alves, and P. Pascoal-Faria, “JANUS: an open-source 3D printable perfusion bioreactor and numerical model-based design strategy for tissue engineering,” Frontiers in Bioengineering and Biotechnology, vol. 11, Dec. 2023. doi: 10.3389/fbioe.2023.1308096

[10] J. P. Efromson, S. Li, and M. D. Lynch, “Biosamplr: An open source, low cost automated sampling system for bioreactors,” HardwareX, vol. 9, p. e00177, Apr. 2021. doi: 10.1016/j.ohx.2021.e00177

[11] A. Jönsson, A. Toppi, and M. Dufva, “The FAST pump, a low-cost, easy to fabricate, SLA-3d-printed peristaltic pump for multi-channel systems in any lab,” HardwareX, vol. 8, p. e00115, oct 2020. doi: 10.1016/j.ohx.2020.e00115

[12] N. Cohen, E. Sicher, C. Ayala-Garcia, I. M. Sanchez-Fayos, L. Conterno, and S. Ugur Yavuz, “Innocell bioreactor: An open-source development to produce biomaterials for food and packaging based on fermentation processes,” Fermentation, vol. 9, no. 10, p. 915, Oct. 2023. doi: 10.3390/fermentation9100915

[13] J. Haseloff, “Biomaker.” [Online]. Available: https://www.biomaker.org/

[14] C. Davidson-Pilon, “Pioreactor.” [Online]. Available: https://pioreactor.com/

[15] L. Parret, K. Simoens, B. Horemans, J. De Vrieze, and I. Smets, “Preprint: Establishing a co-culture aggregate of N-cycle bacteria to elucidate flocculation in biological wastewater treatment,” bioRxiv, 2024. doi: 10.1101/2024.07.02.601720. [Online]. Available: https://www.biorxiv.org/content/early/2024/07/02/2024.07.02.601720

